# A proteome-wide screen for membrane-interactions in intrinsically disordered regions of transmembrane proteins reveals a role in disease

**DOI:** 10.1101/2025.08.15.670465

**Authors:** Joscha Rombach, Tommas Theiss Ehler Nielsen, Jan Hendrik Schmidt, Junior Agenant, Andreas Haahr Larsen, Kenneth Lindegaard Madsen

**Affiliations:** Department of Neuroscience, Faculty of Health and Medical Sciences, Panum Institute, University of Copenhagen, Blegdamsvej 3b, 2200 Copenhagen, Denmark

**Keywords:** intrinsically disordered regions, transmembrane proteins, membrane interaction, protein-lipid interactions, amphipathic helices, protein trafficking, CPP, cell-penetrating peptide, peptide array, liposomes, database

## Abstract

Transmembrane proteins mediate essential cellular processes including signaling, transport, and ion flux. Besides their well-characterized structured domains, most contain intrinsically disordered regions, whose biological roles remain poorly understood. Evidence suggests that the functions of intrinsically disordered regions are context-dependent, a trait particularly relevant when anchored to cellular membranes. In this study, we probed peptide arrays with fluorescent liposomes to generate a high-resolution, proteome-wide map of membrane-interaction sites within intrinsically disordered regions of human transmembrane proteins. Screening 4,000 proteins, we identified membrane-interaction sites in ∼60% of cases. Among these, ∼63% represent amphipathic helices, while ∼17% resemble cationic cell-penetrating peptides. We demonstrate that membrane-interaction motifs can influence subcellular localization and may contribute to both physiological and pathological processes. Our findings establish membrane association as a key functional aspect of intrinsically disordered regions and provide a valuable resource for discovering non-canonical regulatory mechanisms in transmembrane proteins. The resource is available at [MemRIDRdb](https://joscharombach.github.io/MemRIDR/).

## Introduction

Transmembrane proteins (TMPs) account for ∼25% of proteins encoded in the human genome. They carry out key functions including receptor-, transporter- and channel-activity and are targets for more than 60% of marketed drugs^1,2^. Structural studies, combined with advanced algorithms like AlphaFold, are rapidly elucidating the molecular mechanisms underlying the core functions of TMPs. However, it is becoming evident that unstructured intrinsically disordered regions (IDRs) within TMPs equally play functional roles^3^. Along with IDRs being more prevalent in TMPs than in soluble proteins, this indicates that IDRs may play a particularly significant role in TMP function^4^.

IDRs impact a wide range of biological processes and diseases, yet our understanding of their function remains limited^3^. IDR function is highly dependent on the physicochemical environment, and in TMPs, IDRs are naturally situated in a membrane-proximal context. To date, around 29 IDRs in human TMPs have been found to interact with membranes^5^, serving diverse functions, including controlling protein-protein interactions, shaping membranes, linking compartments, and regulating protein function and trafficking. Prominent examples of functional membrane interactions include electrostatic interactions in stargazin^6^, synaptotagmin-1^7^, VAMP2^8^, Kir2.2^9^, Ssy1^10^, as well as insertion of transient structural motifs such as the amphipathic helices (AHs) of Ist2^11^, peripherin-2/rds^12^, REEPs^13^, Reticulons^14^, IFITMs^15^, and FAM134B^16^.

In this study, we mapped membrane binding sites in IDRs from TMPs across the entire human proteome, identifying general motifs that regulate their function. Using peptide arrays and a custom analysis pipeline, we probed the liposome-interaction of all intrinsically disordered cytosolic termini of 4000 human TMPs, finding 2974 significant membrane binding sites. Computational analysis revealed that AHs constituted the majority (∼63%), while cationic amino acid patches made up ∼17% of sites and the remaining 20% lacked identifiable common biochemical features. By further analysis, we identified AHs in the short cytoplasmic tails of glycosyltransferases and demonstrated that these confer localization to the Golgi apparatus. AHs can also help stabilize membrane curvature in TMPs such as membrane shaping adapter proteins, and here we confirmed such known AHs and identified new AHs in most members of the superfamily. We also highlight the existence of AHs in IDRs of channels and transporters and draw parallels to the amphipathic helix 8 of GPCRs. Finally, we show that AHs are under significant evolutionary constraint and associated with a disproportionate number of pathological single nucleotide polymorphisms (SNPs). Ultimately, we present a map of membrane binding in IDRs of TMPs, which includes known membrane binding regions and significantly increases their number, allowing for the identification of new and unconventional mechanisms that regulate TMP function. The map is accessible as the Membrane Recruited IDRs database, [MemRIDRdb](https://joscharombach.github.io/MemRIDR/).

## Results

### A high-density peptide array to map membrane binding sites within IDRs

Peptide arrays allow for simultaneous probing of thousands of peptides and can be used to characterize interactions down to the amino acid level^17^. To screen for protein-membrane interactions in IDRs of human TMPs, we extracted all cytosolic, membrane proximal C- and N-terminal IDRs from TMPs in the human proteome following exclusion of sequences with annotated domain structure (4685 termini) (Figure 1A, materials and methods for details). Following the release of AlphaFold, regions with predicted domain structure were excluded post hoc (Figure S1, materials and methods for details). To allow for precise mapping of binding sites, we generated overlapping 16-mer peptides from each IDR at dual amino acid resolution (Figure 1A). The 203,031 resulting peptides were synthesized in randomized 20×20 *µ*m^2^ fields on a functionalized glass slide separated by 10-µm spacers. 18,465 controls were added, including empty fields and the membrane-binding peptide Hecate, as negative and positive controls, respectively. Controls for the contribution of charge, hydrophobicity, amphipathicity, and helicity to membrane binding were also included (Table S1). The array was probed for membrane binding capacity of individual peptides using liposomes labeled with the carbocyanine dye DiD (Figure 1B, Figure S2A-D). Inspection of fluorescence images showed good separation of signal between fields within the array (Figure 1B), and we did not find indications of signal overspill in surrounding fields (Figure S2E). Importantly, incubation with DiD alone did not result in fluorescence (not shown), demonstrating that we specifically assessed peptide-liposome interactions.

**Figure 1.**
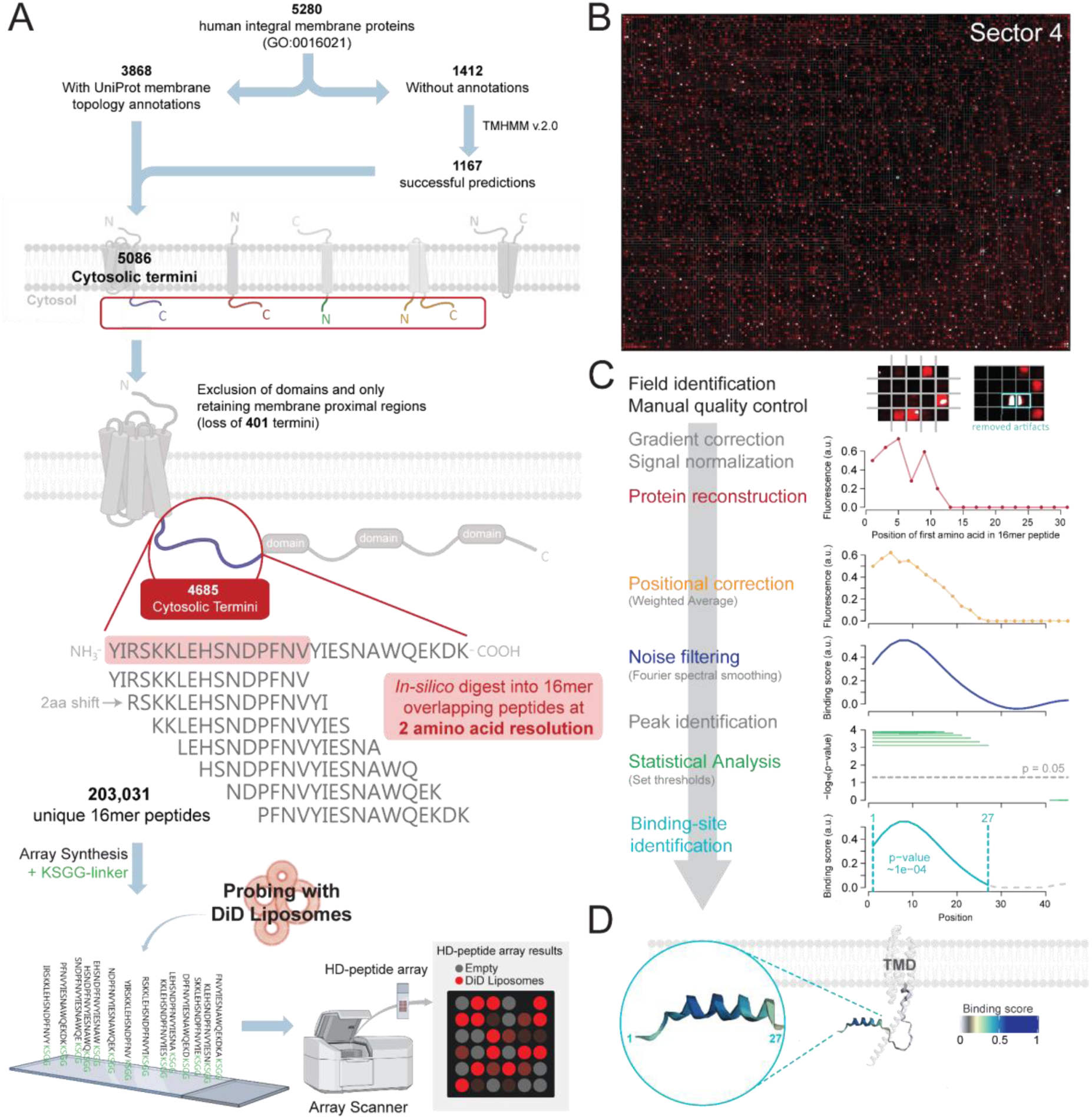
A high-density peptide array to study protein-membrane interactions. **(A)** Intrinsically disordered regions in the cytosolic termini of all human transmembrane proteins (GO:0016021) were extracted and segmented into 16mer overlapping peptides at two amino acid resolution in silico. The resulting 203,031 unique 16mer peptides were synthesized onto an array anchored at the C-termini with a KSGG linker. The peptide array was probed with fluorescent DiD liposomes and imaged using an array scanner. Partly created with Biorender.com. **(B)** Representative image of one of six sectors on the array (sector 4 of replicate 2). The 17 highly fluorescent peptide fields in each corner of the sector contain the membrane binding synthetic amphipathic helix Hecate (Table S1). **(C)** Flow chart illustrates the post-processing pipeline and statistical analysis. Example protein N-terminus of TMEM17 (UniProt accession: Q86X19). **(D)** Binding score mapped onto the N-terminus of the AlphaFold structure of TMEM17 (AF-Q86X19-F1). Binding site highlighted. See also Figure S1-7 and Table S1.

We normalized gradient-corrected fluorescence intensities to the intensity of empty fields and Hecate (Figure S3, see materials and methods), and constructed the membrane binding regions for each IDR using the fluorescence signal from individual peptide fields (Figure 1C, Figure S4A). To correct for peptide position and eliminate noise, we calculated a smoothed weighted-average fluorescence intensity score (hereinafter binding score) for amino acids in each IDR (Figure 1C, Figure S4B-F). To statistically assess the significance of membrane binding peaks within the large dataset, we first created background distributions of the maximum scores expected for peaks of different lengths while considering the actual length of the IDR (example for the longest IDR, see Figure S5B-E). We did this by sampling random fluorescence intensities from peptide fields on the array, assembling them into mock sequences that represent the lengths of all true TMP IDRs, and calculating their binding scores. Next, we identified membrane binding peaks in actual IDRs (Figure S6A and D) and tested their significance by calculating the probability of observing the peaks in the randomly sampled background distributions (Figure S6B-C and E-F). A significance criterion of 0.05 was used to classify peaks as binding and 0.95 for non-binding. A score corresponding to the area under the curve was assigned to each peak (hereinafter AUC score).

### The peptide array offers reproducible and robust screening for membrane binding sites in IDRs of all human TMPs

To evaluate the robustness of the setup, we repeated the experiment on another array with peptides in new random positions and calculated Pearson’s correlation between replicate binding traces for each IDR. Most of the replicate traces were positively correlated (4239 of 4685 traces) (Figure S7A), especially IDRs with significant membrane-binding peaks (2450 of 2494 traces) (Figure S7C). Assessing individual binding scores for amino acid pairs also revealed a strong correlation between the two datasets (Figure S7D, R^2^ 0.8, slope=0.87, p-value < 2.2e-16). Evaluation by the statistical analysis showed that most of the identified binding sites overlapped by more than 95% (Figure S7E-G), and 2247 binding sites were identified in both replicates (min. 50% overlap) (Figure S7H). Finally, the AUC scores for overlapping binding sites were correlated (R^2^ 0.91, slope=0.938, p-value < 2.2e-16) (Figure S7I). AUC scores for overlapping sites were averaged and Fisher’s method was used to combine the p-values from the two independent sets.

### More than half of TMP IDRs across structural classes contain membrane binding regions

Assessing prevalence, we identified 2257 TMPs (∼56%) that contain significant membrane binding sites (Figure 2A). The location and patterns of membrane binding are strikingly similar for many TMPs, mostly located proximal to the transmembrane segments (Figure 2A and B), without any distinct preference for N or C-termini (2000 sites in 3373 C-termini = 59 %; 974 sites in 1428 N-termini = 0.68%). 475 TMPs contain two or more binding sites. The lengths of binding sites range from ∼10 to 80 amino acids (Figure 2C). Notably the lengths are not normally distributed but show two apparent peaks at 20 and 30 amino acids. Examples of TMPs with different binding site lengths are shown in Figure 2D.

**Figure 2.**
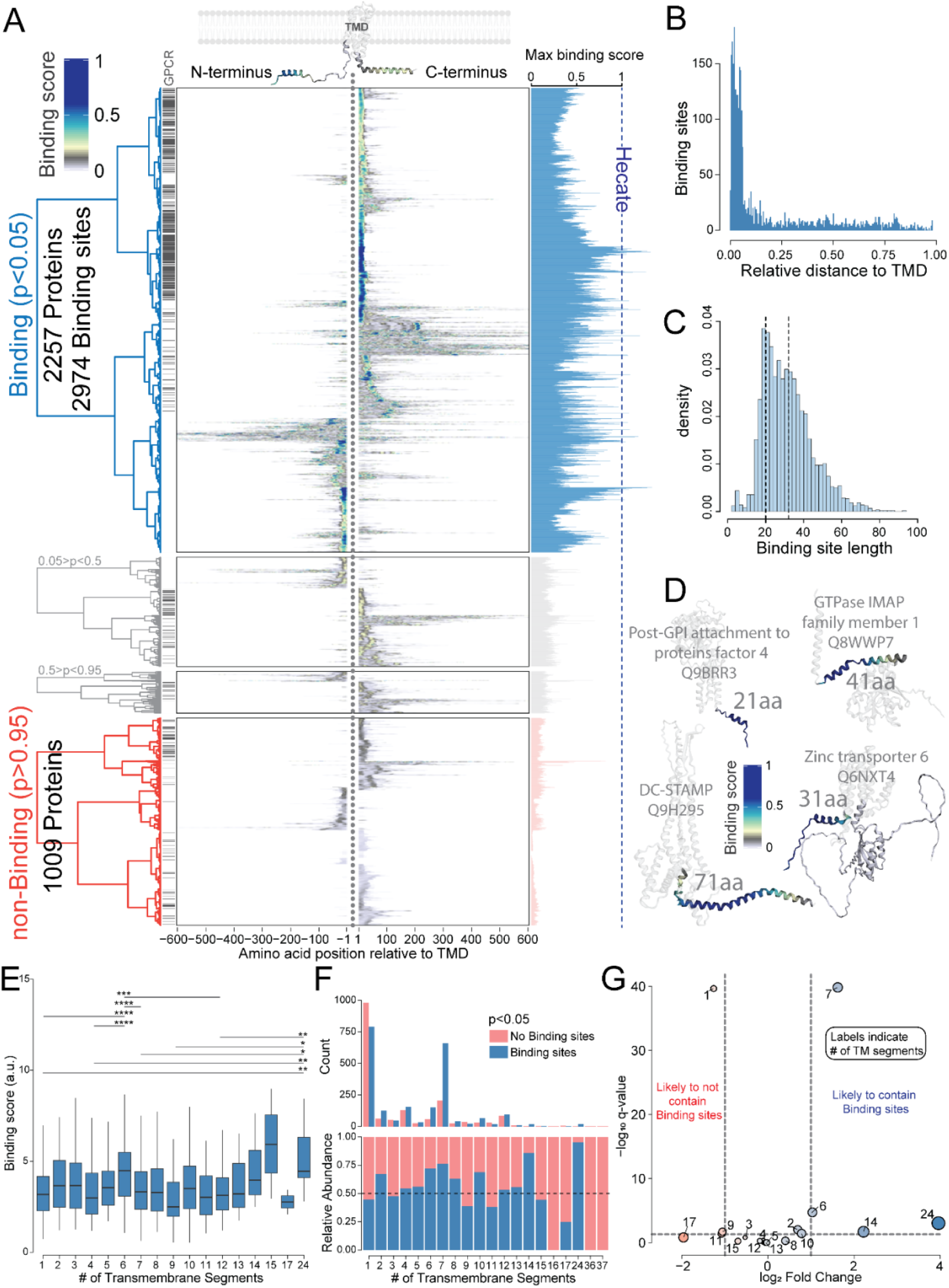
More than half of TMP IDRs across structural classes contain membrane binding regions. **(A)** Heatmaps showing the binding scores (i.e. membrane binding propensity) in the termini of all transmembrane proteins on the array with liposome binding sites (top/blue) and without liposome binding sites (bottom/red). Amino acid positions are plotted relative to the transmembrane domain (TMD) shown in the middle with negative values indicating the distance from the TMD within the N-terminus and positive values indicating the distance within the C-terminus. The barplot for each row (right) shows the maximum binding score in each protein. Dashed blue line indicates the average binding score for the positive control Hecate. The rows of the heatmaps were clustered based on their binding profiles using Ward’s hierarchical agglomerative clustering. The discrete grid-like annotation at the tip of the dendrogram highlights G protein coupled receptors (GPCRs). **(B)** Histogram showing the relative distance of binding sites from the TMD. **(C)** Histogram (with bin width 2 amino acid), showing the distribution of binding site lengths. **(D)** Binding scores of binding sites mapped onto AlphaFold structures of four TMPs with binding site lengths falling within the three normal distributions of the fit in panel C. Generated with [MemRIDRdb](https://joscharombach.github.io/MemRIDR/). **(E)** Box-and-whiskers showing the distribution of binding strength (AUC scores) of sites for TMPs grouped by topology. Error bars show the 5-95 percentile. Pairwise comparison is a non-parametric Kruskal-Wallis test with Holm’s correction for multiple comparison. ****p-value ≤ 0.0001, ***p-value ≤ 0.001, **p-value ≤ 0.01, *p-value ≤ 0.05, comparisons not shown p-value > 0.05. **(F)** Histograms showing the absolute (top) and relative (bottom) abundance of binding sites in TMPs with different topologies. **(G)** Enrichment analysis comparing the abundance of binding site-containing TMPs in each topology group with the ratio for all TMPs. Multiple Fisher test p-values corrected using the false discovery rate (q-value). See also Figure S8-S9.

To investigate whether binding sites are specific to TMPs with distinct architectures, we grouped TMPs by topology (Figure S8). While most groups display generally similar binding site scores (AUC), only 6TM and 24TM show a significant increase (Figure 2E). However, the proportion of binding sites in each group varies widely (Figure 2F, bottom). In absolute numbers, the 1TM group contains the most binding sites, followed by 7TM, represented mainly by helix 8 in GPCRs (Figure 2F, top). Enrichment analysis revealed that 1TM and 9TM topologies are significantly depleted, while 6TM, 7TM, 14TM, and 24TM are significantly enriched in binding sites (Figure 2G) alongside their binding scores.

### Enrichment of post-translational modifications highlights membrane-binding sites as regulatory hotspots

To investigate whether binding sites are regulated by specific post-translational modification, we tested their enrichment in binding sites against non-binding sites (Figure S9A). S-palmitoylation was increased 4-fold in binding sites, while methylation and ubiquitination were also significantly higher, by ∼2-fold. Although significantly enriched, phosphorylation occurs at comparable frequencies in both groups. Notably, S-palmitoylation has been shown to drive membrane binding in concert with membrane binding motifs^18,19^, which have been shown to direct and confer palmitoylation^20,21^. To assess whether post-translational modifications have a specific distribution pattern in binding sites, we plotted the probability density of modified sites with respect to the center of binding sites (Figure S9B). This revealed that S-palmitoylation and methylation are more probable N-terminally in binding sites. Ubiquitination and phosphorylation are more evenly distributed throughout the binding sites, although ubiquitination exhibits a small C-terminal peak.

### Biochemical analysis of membrane binding reveals amphipathic helices as the most prominent binding mode

Comparison of general amino acid properties within binding and non-binding regions did not reveal any difference in terms of hydrophobicity (Figure 3A). However, while the net charge of non-binding regions showed normal distribution around 0, binding regions exhibited a right shift with a median net charge of +1 (Figure 3B) driven by overrepresentation of positively charged residues (K and R), as expected for sequences binding to net-negatively charged liposomes. Although net hydrophobicity did not differ from non-binding regions, hydrophobic amino acids were clearly enriched in binding sites, especially aromatic amino acids (Figure 3C). Notably, this pattern was not affected by the membrane binding amphipathic helix 8 in GPCRs (647 binding sites) (Figure S10A).

**Figure 3.**
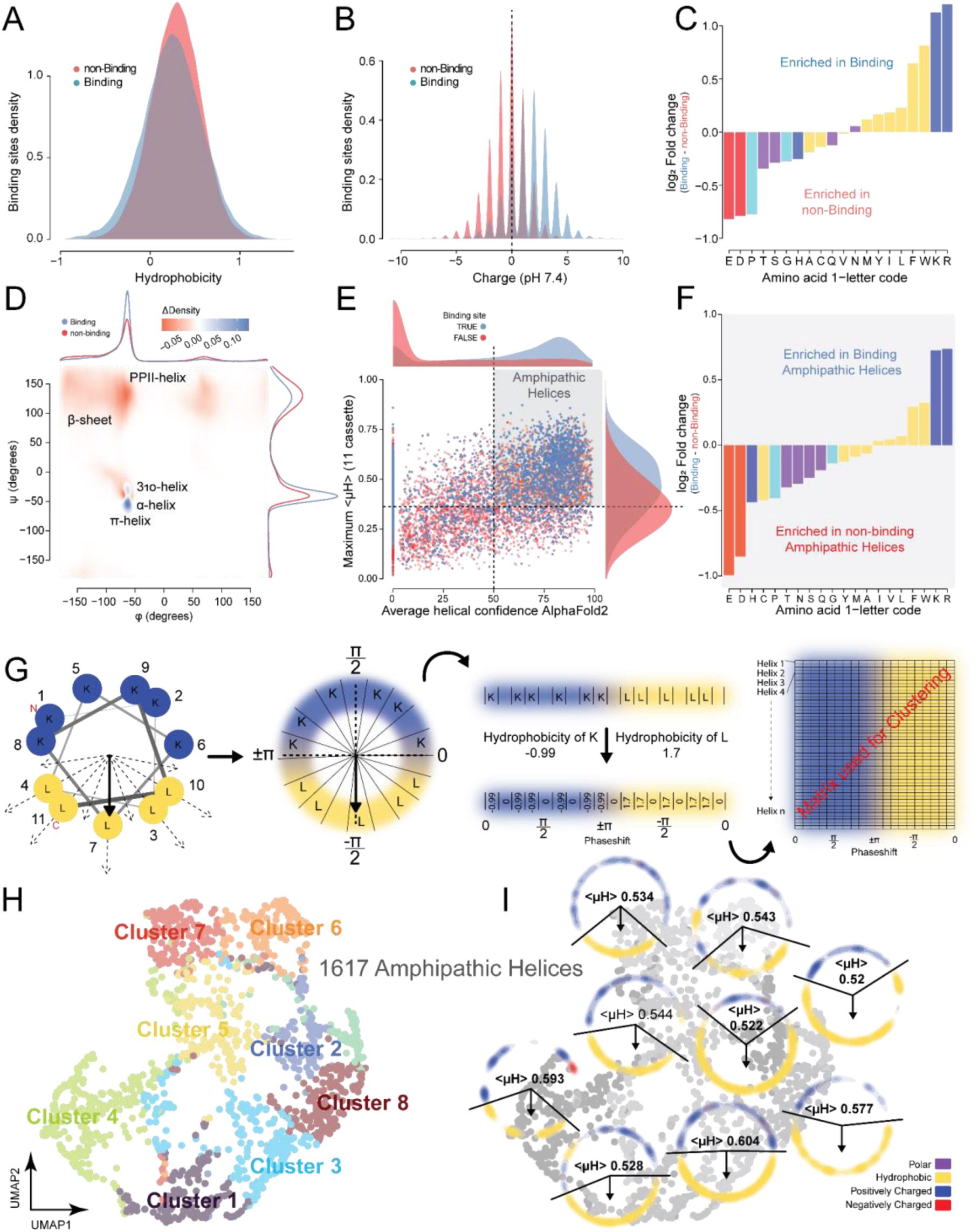
Biochemical analysis of membrane binding reveals amphipathic helices as the most prominent binding mode. **(A)** Density plots showing the mean Hydrophobicity in binding sites (blue) and in nonbinding regions (red). **(B)** Density plots showing the net charge in binding sites (blue) and in nonbinding regions (red) at pH 7.4. **(C)** Amino acid enrichment in binding sites vs non-binding sites. **(D)** Ramachandran plot showing the difference between peptide bond φ and ψ angles within and outside of binding sites. Density difference reported as log2 fold change between the presence of the respective structural elements in the binding and non-binding groups. Marginal plots show density. **(E)** 2D point and density plot showing cutoffs used to classify regions as AHs. Vertical dashed line at 0.36 is the median <µH> of the 11-mers with the maximum <µH> from all non-binding sequences. **(F)** Enrichment of amino acids in AHs that bind liposomes relative to those that do not. **(G)** Schematic illustrating how AHs were encoded for clustering in panel H (see also Method section) **(H)** UMAP of hydrophobic profiles of helical wheel projections of binding sites qualifying as AHs. Clustered by K-means. **(I)** Helical wheel density plots for each cluster in G. Yellow, density of hydrophobic amino acids W, F, Y, I, L, M, C; Blue, density of positively charged amino acids K and R; Purple, density of S and T; Green, density of P; Red, density of E and D; <µH>, mean hydrophobic moment. See also Figure S10-12.

The combined enrichment of hydrophobic and positively charged residues suggest that a binding is not simply driven by positive charge, but likely also by amphipathic structural motifs, e.g. in the form of AHs. To evaluate the proportion of binding sites representing putative AHs, we compared AlphaFold predicted backbone *ϕ* and *ψ* angles between binding and non-binding regions. This analysis confirms that ɑ-helical structures are enriched in membrane binding sites (Figure 3D). Furthermore, the mean hydrophobic moment (<µH>) was significantly higher when calculated for binding sites than for non-binding sites, indicating that the ɑ-helical structures in binding sites display amphipathic properties (Figure 3E). To extract a subset of membrane binding sequences that can be categorized as AHs, we set a threshold of at least 50% predicted helical structure (as determined by AlphaFold) and a <µH> higher than the median moment (<µH> > 0.36) observed for non-binding sequences (Figure 3E). Remarkably, ∼63% of all binding sequences can be classified as AHs at the set threshold. Although AHs appear to make up a large share of all binding sites, we noted that several non-binding sites exhibited similar properties when evaluating <µH> and helicity. However, the proportion of AHs in binding sites was significantly higher than in non-binding regions (Fisher’s exact test; odds ratio 4.8, 95% CI 4.4-5.3, p-value < 2.2e-16).

To determine the properties that define membrane binding AHs, we compared the amino acid composition to non-binding AHs from the same quadrant all qualified by <µH> and helical propensity. To avoid bias due to high sequence similarity of helix 8 in olfactory GPCRs, we removed these from the analysis. We observed an enrichment of positively charged and aromatic residues (F and W) in binding AHs, while negatively charged amino acids, as well as histidine, cysteine, proline, and polar (T, N, S, Q) residues were enriched in non-binding AHs (Figure 3F). To evaluate the distribution of amino acids along the binding and non-binding AHs, we created helical density maps based on helical wheel projections (Figure S11A). Interestingly, we noted that amino acids in binding and non-binding AHs exist in different phases in the sense that the non-binding helices showed a highly localized density map with first and last residue in the hydrophobic face suggesting that this configuration compromises membrane binding (Figure S11B). Binding AHs also exhibit higher hydrophobic density (Figure S11C) driven strongly by aromatic residues (F and W) concentrated in the center of the hydrophobic face (Figure S11D). In the hydrophilic face, binding AHs have a high positive charge density, while negative charges are generally excluded (Figure S12A-B).

To assess liposome binding of peptides with extreme variations in charge, hydrophobicity, and helicity, we designed several controls and examined their binding to liposomes in the array setup (Table S1, Figure S10B-C). Although highly positively charged peptides readily bound to liposomes, charge alone did not determine the degree of binding (Figure S10C-D). Keeping charge constant but ordering amino acids to represent an amphipathic helix (Hecate) significantly increased the level of membrane binding compared to a shuffled version of the same peptide (A-shuffle). Substitution of alanines in A-shuffle to glycines (G-shuffle), adding more degrees of rotational freedom to the peptide backbone, decreased membrane binding even further, while substitution of positive charges for glutamate (Hecate K→E) completely abolished membrane binding. We also detected weak binding to the hydrophobic membrane-embedded motif of GPAT4, liveDrop.

### Amphipathic helices in TMPs may sense a wide range of membrane curvatures

AHs are thought to sense or induce different membrane curvatures depending on the properties of the hydrophobic face, as well as their hydrophobicity/insertion depth in the membrane^22^. Specifically, molecular dynamics simulations have suggested that AHs with wide hydrophobic faces can sense negative membrane curvature and that there is a correlation between the AH insertion depth in the membrane and its preferred mean curvature^23^. To address what types of AHs exist in TMPs, we clustered all putative AHs based on their 2D helical wheel hydrophobicity profiles (Figure 3G-H). This revealed a highly diverse nature of the hydrophobic faces, some as narrow as 90° to highly hydrophobic AHs with faces spanning up to 270° (Figure 3I). This implies that AHs might serve diverse functions in TMPs, including targeting TMPs to membranes of variable curvatures and partaking in shaping membranes to specific curvatures.

### Glycosyltransferases navigate membranes by means of amphipathic helices

We recently demonstrated that amphipathic motifs drive endocytosis of TMPs but also suggested that they confer association with/retention to the biosynthetic pathway^24^. To explore the functional role, we investigated the putative differential impact of membrane-binding AHs on subcellular localization within the simplest topological class of TMPs, the 1TM proteins. Analyzing Gene Ontology (GO) term enrichment for AH-containing 1TM proteins revealed that cell surface, cell adhesion and protein kinase related 1TMs were significantly underrepresented, while glycosyltransferases emerged as the major 1TM group enriched for membrane binding AHs (Figure 4A). Extraction of all annotated glycosyltransferases in the PANTHER database revealed a single membrane binding region in the short tails of most glycosyltransferases (Figure 4B). Enrichment analysis evaluating the sub-cellular localization using the COMPARTMENTS database^25^ or GO cellular components, revealed that AH-containing glycosyltransferases are enriched in the Golgi apparatus rather than in the ER, the general localization for glycosyltransferases (Figure 4C). To experimentally test whether such AHs in glycosyltransferases can drive their localization to the Golgi apparatus, we extracted the AH from Golgi-localized WSCD2^26^ and the membrane binding region from ER-localized UGT1A4 as control^27^ (Figure 4D) and fused them to the C-terminus of the trafficking inert interleukin-2 receptor subunit *α* (Tac). Consistent with our previous findings, the AH from WSCD2 drove efficient endocytosis of Tac, while the non-amphipathic binding site from ER-resident UGT1A did not (Figure 4E-F). Moreover, the Golgi-resident WSCD2 AH increased the Golgi localization of Tac, while the charged sequence from UGT1A did not (Figure 4G-H). This suggests that AHs in glycosyltransferases act as facilitators of ER exit and forward trafficking leading to increased steady-state localization to the Golgi apparatus; a hypothesis supported by mutational studies of the amphipathic helix 8 in GPCRs, where AH disruption leads to GPCR accumulation in the ER^28–31^.

**Figure 4.**
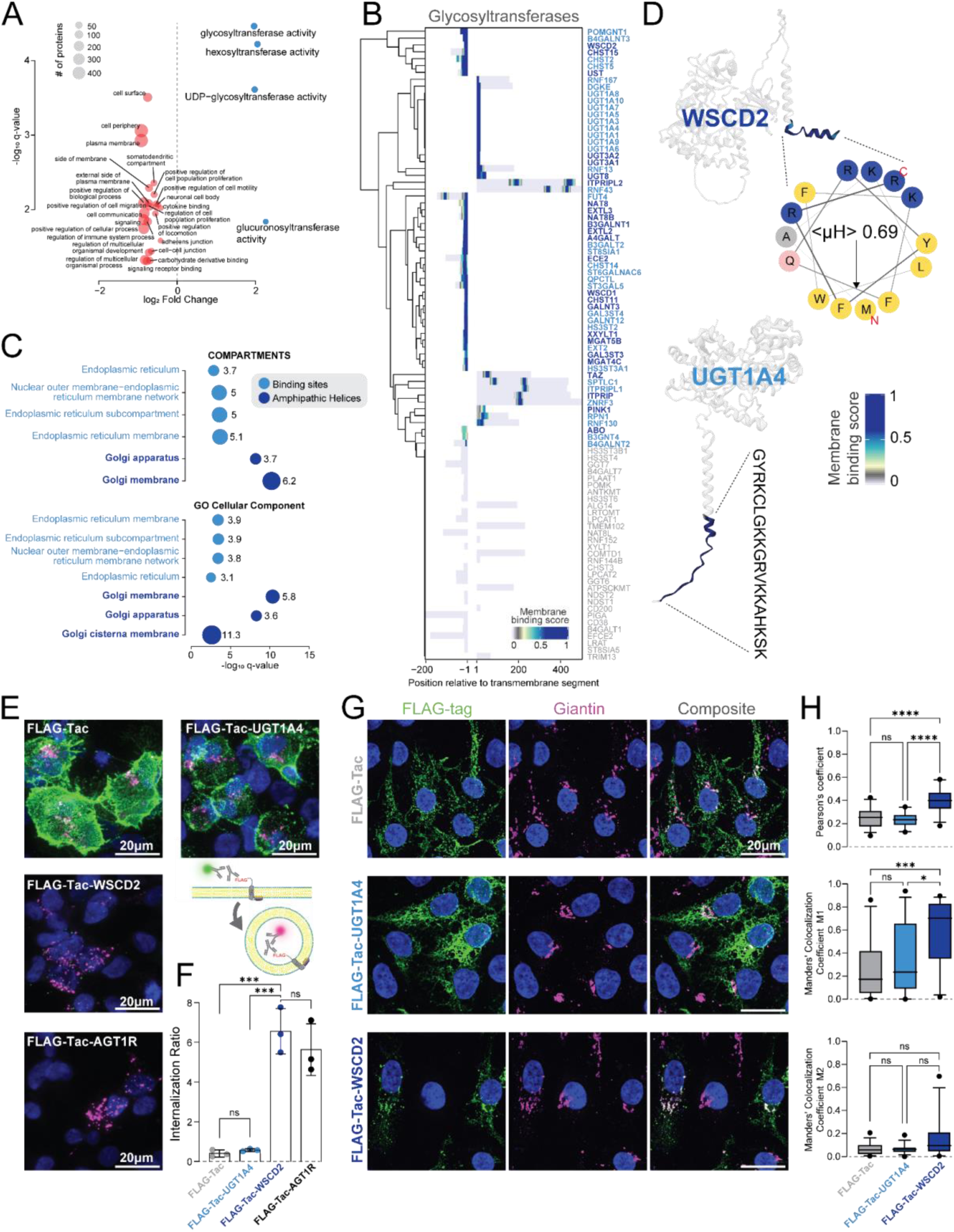
Glycosyltransferases navigate membranes by means of amphipathic helices. **(A)** Overrepresentation analysis of PANTHER ontology terms associated with membrane binding and non-binding for 1TM proteins. FDR corrected p-value, significance threshold 0.05. **(B)** Heatmap showing the binding scores for all 1TM glycosyltransferases extracted using PANTHER protein class annotations. The rows of the heatmap were clustered based on the Euclidean distance between binding profiles using the complete linkage hierarchical clustering algorithm. Protein names in dark blue; with AHs, light blue; with binding sites, grey; no binding sites. **(C)** Overrepresentation analysis for glycosyltransferases containing AHs and non-AH binding sites using either the COMPARTMENTS database^25^ or GO cellular compartments extracted from the PANTHER database. **(D)** Binding scores mapped onto the AlphaFold structures of Golgi localized glycosyltransferase WSCD2 and ER localized UGT1A4. Generated with [MemRIDRdb](https://joscharombach.github.io/MemRIDR/). The helical wheel projection and mean hydrophobic moment (<*µ*H>) of the WSCD2 AH is shown. **(E)** Representative summed projections of z-stack images showing immuno-staining for FLAG-Tac (Tac), FLAG-Tac-UGT1A4, FLAG-Tac-WSCD2 and the positive control, FLAG-Tac-AGT1R following 30 min of internalization in the presence of recycling inhibitor monensin. Magenta; internalized, green; surface, blue; DAPI. Scale bar 20 µm. **(F)** Medians of three independent internalization experiments with >24 cells per construct per experiment. Error bars; SD. Pairwise comparison; one-way ANOVA with Tukey’s multiple comparison test. ***p-value ≤ 0.001, ns > 0.05. **(G)** Representative confocal images (single z-plane) of colocalization between Golgi-marker Giantin (magenta) and FLAG-stain (green) for FLAG-Tac, FLAG-Tac-UGT1A4 and FLAG-Tac-WSCD2. DAPI in blue. Scale bar 20 μm. **(H)** Box-and-whiskers of Pearson’s correlation coefficient and Manders’ colocalization coefficients M1 and M2 for colocalization between Giantin and FLAG-Tac-UGT1A4, FLAG-Tac-WSCD2 and FLAG-Tac as a reference. M1 represents the fraction of Giantin (magenta) that contains FLAG-tag (green), while M2 is the fraction of FLAG-tag (green) containing Giantin (magenta). Error bars show the 5-95 percentile and points show outliers. At least 27 cells per condition. The pairwise comparison is a non-parametric Kruskal-Wallis test with Dunn’s correction for multiple comparison. ****p-value ≤ 0.0001, ***p-value ≤ 0.001, **p-value ≤ 0.01, *p-value ≤ 0.05, ns > 0.05. See also Figure S13-15

### Amphipathic helices are ubiquitous in membrane-shaping adapter proteins

AHs are already recognized to serve a role in shaping cellular membranes in the membrane-shaping adaptor protein (MSAP) superfamily^32^. MSAPs shape membranes by sensing and inducing curvature and act as adapters for protein-protein interactions, e.g. during cargo trafficking or during fission and fusion of mitochondria or peroxisomes^33–38^. The most studied families include reticulons, receptor expression enhancing proteins (REEPs) and atlastins^39^. To qualify our data and identify additional AHs in the MSAP superfamily, we extracted MSAPs and evaluated their binding sites (Figure S13A). We identified membrane binding AHs in most cytosolic termini, including all previously identified AHs (Figure S13A+C and S14). The AHs exist in distinct clusters with hydrophobic faces as narrow as 45° and some wider than 180° (Figure S13B), suggesting that MSAP AHs can be divided into sensors of different curvatures or with distinct functions^23^. We encountered AHs in several peroxisomal TMPs (Figure S15), including the known curvature-generating AH in PEX11A/HsPEX11p (H3) involved in membrane elongation during peroxisome division^34,35^, a second membrane binding AH in PEX11A and an AH in the N-terminus of PEX11G. We also identify an AH in the PEX11A interaction partners Mitochondrial Fission factor (MFF) and FIS1, which together orchestrate peroxisome division^40,41^, as well as putative novel AHs in mitochondrial fusion proteins mitofusins (MFNs). MFNs are known to contain AHs in the HR1 and HR2 domains, and in the C-terminal TM-proximal region, which facilitate mitochondrial targeting and fusion by MFN1^36–38^. We confirmed the existence of membrane-binding AHs in the TM proximal regions of MFN1 and expanded these findings to MFN2. An unconventional N-terminal TM proximal AH has also been reported in MFNs^37^. However, we could not verify membrane binding for the reported N-terminal AH and instead identified a classical AH in an adjacent region. We further identify AHs in mitochondrial proteins such as fusion proteins MID51 and MID49, as well as import receptor subunits TOM7, TOM20 and TOM22 (see MemRIDRdb). Ultimately, AHs appear to be prominent features in most MSAPs and might be involved in a wide range of membrane modulatory processes, including curvature stabilization, protein trafficking, and membrane fission and fusion.

### Amphipathic helices in channels and transporters resemble helix 8 in GPCRs

To investigate how AHs are distributed between functional groups of TMPs, we performed a functional enrichment analysis to identify significantly overrepresented GO terms associated with TMPs containing putative AHs. This revealed that TMPs associated with GPCR activity, channel activity, and transporter activity are enriched for AHs (Figure 5A, Table S1). Membrane binding in the C-termini of GPCRs coincides with its amphipathic helix 8 (Figure 5B-C), a known membrane binding motif ^24,42–44^. In particular, the analysis highlights olfactory receptors, with the vast majority containing a membrane binding helix 8 in their short cytosolic C-termini (Figure 5D). Extracting the 153 channels and 176 transporters annotated in the PANTHER database and assessing the hydrophobicity profiles of their AHs, revealed profiles similar to those of GPCRs (Figure 5B). Clustering the termini according to the location of the binding site, we noted that the distribution of binding sites within the cytosolic termini of most channels and transporters is highly reminiscent of the profiles found in GPCRs although with a trend towards more membrane distal localization of the binding sites in a subset of the channels and transporters (Figure 5D, with structural examples in 5E). Helix 8 of GPCRs have a wide variety of functions ranging from structural integrity^29,45^ to mechanosensing^46^, signaling^47,48^ or trafficking^24,31,49^. Consequently, one might speculate that AHs in channels and transporters may play similar roles.

**Figure 5.**
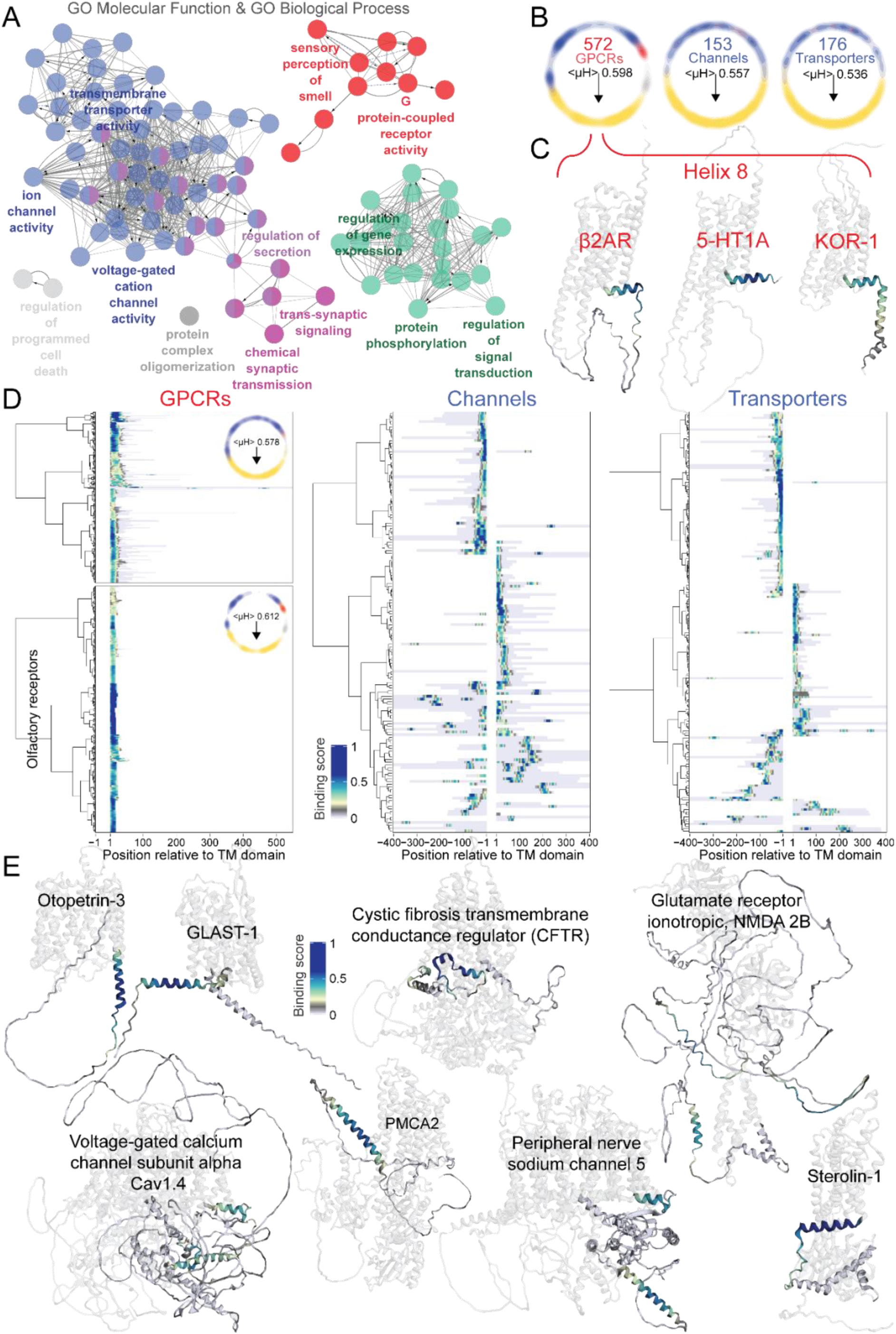
Amphipathic helices in channels and transporters resemble helix 8 in GPCRs. **(A)** ClueGO functional analysis of AH containing TMPs using GO terms biological process and molecular function. Group labels were selected based on the smallest p-value and number of associated genes. Arrows indicate ontology relations. See Table S2 for full table and details. **(B)** Average helical wheel density plots for AHs in GPCRs, channels and transporters. **(C)** Binding scores mapped onto AlphaFold structures of GPCRs, showing the mapping of binding sites to helix 8. Generated with MemRIDRdb. **(D)** Heatmaps showing Binding scores for AHs in the termini of *left:* GPCRs (Inset shows heatmap of binding scores for all olfactory receptors), *middle:* Channels, *right:* Transporters. **(E)** Binding scores of AHs mapped onto AlphaFold structures of selected transporters and channels. Generated with MemRIDRdb.

### Pathological single nucleotide polymorphisms (SNPs) disrupt evolutionarily constrained amphipathic helices

To gauge the evolutionary conservation of the identified binding sites, we extracted all orthologs for each TMP from the Ensemble resource and calculated the relative substitution rate for each amino acid from multiple sequence alignments (see materials and methods)^50^. Interestingly, we found that binding sites display higher evolutionary constraint with fewer allowed substitutions than non-binding IDRs (Figure 6A). AHs displayed a particularly high evolutionary constraint, with a median amino acid substitution rate ∼26% lower than for non-binding IDRs. This suggests that the identified binding sites, especially AHs, are structurally or functionally important.

**Figure 6.**
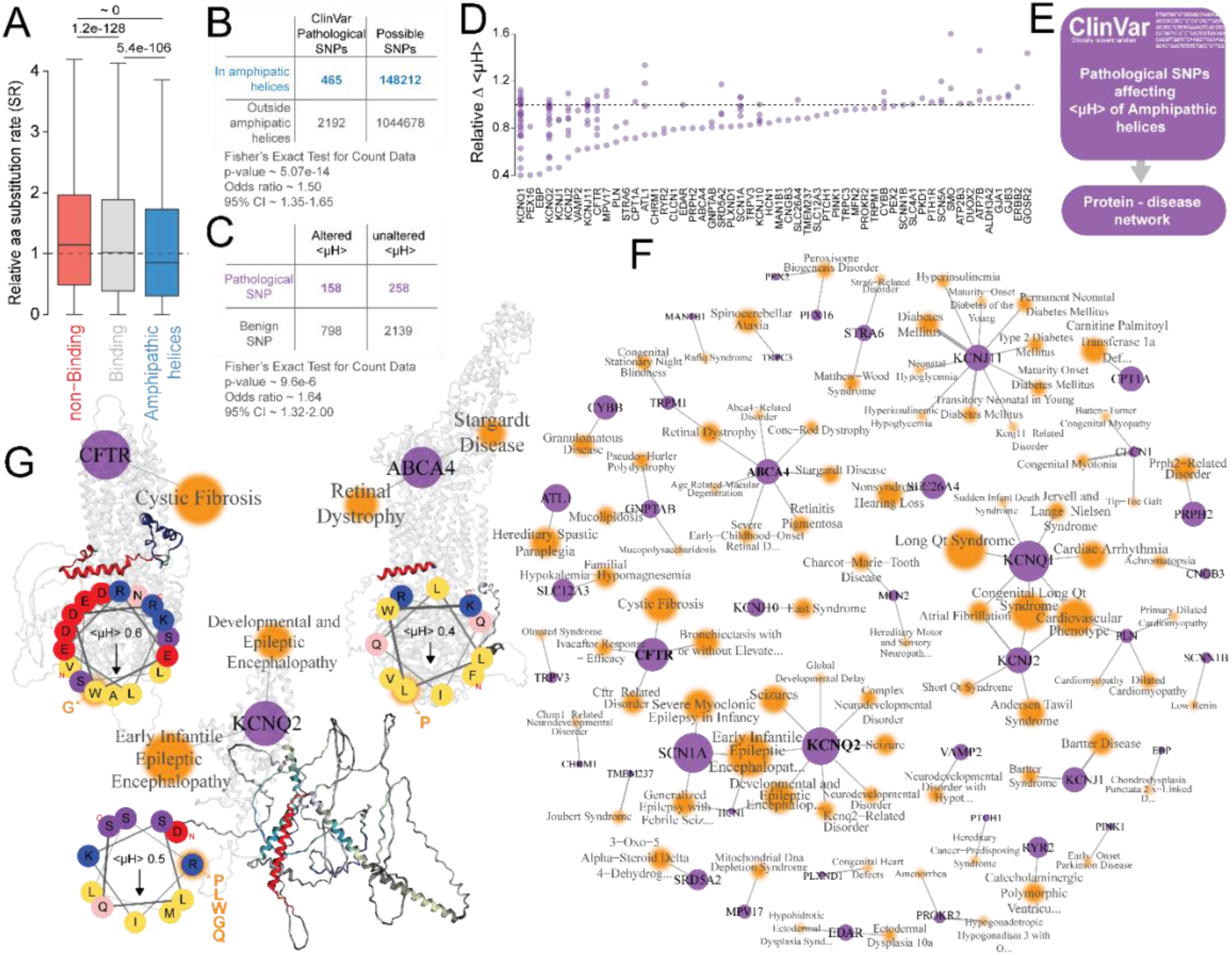
Pathological SNPs disrupt evolutionarily conserved amphipathic helices. **(A)** Box plots (10-90^th^ percentile) show the distribution of relative substitution rates (SR) for amino acids in non-binding regions, binding sites and AHs (relative to the average substitution rate of amino acids in the corresponding protein). Lower SR = higher evolutionary constraint. Pairwise Wilcoxon rank sum test with Bonferroni correction; median of amino acid SR in non-binding regions: 1.15, n=226384; median of amino acid SR in binding sites: 1.02, n=92095 (81 binding sites excluded due to missing orthologs); median of amino acid SR in AHs: 0.85, n=51529. non-binding regions vs binding sites, U-statistic 9855930318, p-value 1.20e-128; non-binding regions vs AHs, U-statistic 5104572858, p-value 0; binding sites vs AHs, U-statistic 2207716702, p-value 5.37e-106. **(B)** Contingency table and results of Fisher’s exact test to test whether single nucleotide polymorphisms (SNPs) annotated as pathological in the ClinVar database are equally frequent within and outside AHs. **(C)** Contingency table and results of Fisher’s exact test to test whether the frequency of SNPs that alter the <µH> of AHs is equal for pathological and benign SNPs. **(D)** Plot showing the relative change in <µH> of membrane-binding AHs caused by pathological SNPs with individual protein names. **(E)** Schematic illustrating how the TMP-disease network was created. **(F)** TMP-disease network showing diseases that are associated with SNPs that disrupt the <µH> of membrane-binding AHs in human TMPs. **(G)** Example sub-networks extracted from panel F highlighting the location of amino acid variants in the AlphaFold2 structure of the proteins, and a helical wheel showing how specific pathological variants (orange) affect the membrane-binding AH in the TMP.

To assess their functional significance in relation to disease, we extracted ClinVar^51^ pathological SNPs within AHs. We identified 465 variants in 105 TMPs associated with 300 diseases. There were significantly more pathological SNPs in AHs than expected for IDRs with an odds ratio of ∼1.5 (Figure 6B). Additionally, pathogenic variants in AHs more frequently disrupt the <μH> compared to benign variants (Figure 6C-D). Creating a network of TMPs and diseases associated with <μH>-disrupting SNPs, revealed several clusters, including cystic fibrosis, cardiovascular phenotype, and epilepsy clusters (Figure 6E-G).

### Cationic regions in transmembrane protein IDRs drive membrane binding

Finally, we addressed the binding sites that do not represent AHs. The hydrophobicity of these binding sites is lower than for AHs and the non-binding background (Figure 7A), while the net charge is higher (Figure 7B). By sequence alignment we identified a cluster consisting of 31 highly similar binding sites that originate from clustered protocadherins (Figure 7C). Interestingly, the extreme C-terminal of *γ*-protocadherins is known to bind phospholipids, an interaction regulated by phosphorylation^52^. Here, we confirm this finding and identify equivalent C-terminal binding sites in *α*-protocadherins, non-clustered protocadherins, cadherins, and FAT4 (Figure 7D+E). As described by Keeler et al.^52^, binding probably depends on electrostatic interactions between phospholipid headgroups and a short patch of lysines.

**Figure 7.**
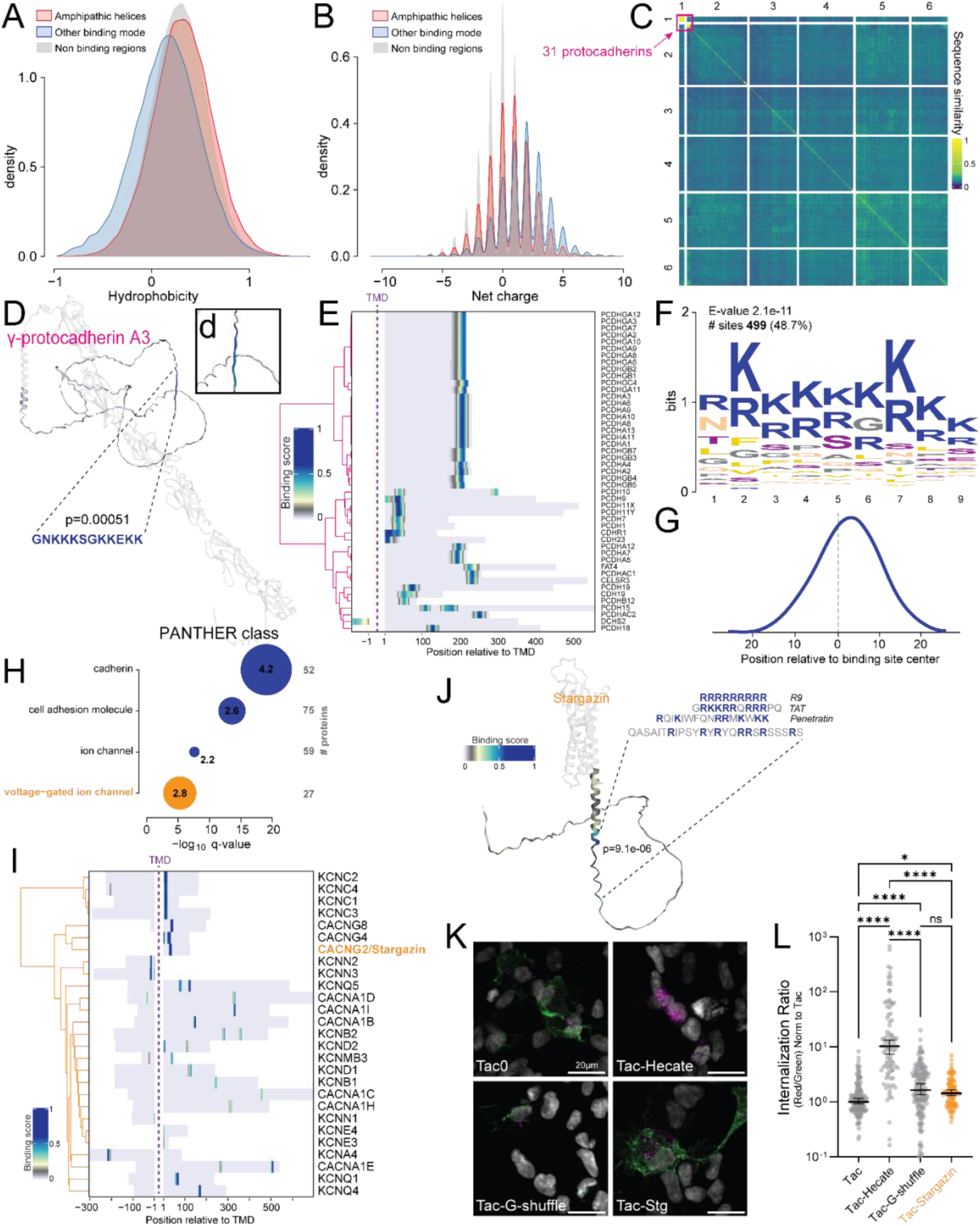
Cationic regions in transmembrane protein IDRs drive membrane binding. **(A)** The distributions of mean hydrophobicity in non-membrane binding regions, membrane binding AHs and other binding sites. **(B)** Density plot showing the net charge calculated at pH 7.4 for the same regions as in A. **(C)** Heatmap showing the sequence similarity between binding sites that do not qualify as AHs. Calculated using the Smith-Waterman algorithm for local alignment. Heatmap rows and columns were clustered using Ward’s hierarchical agglomerative clustering. Pink box highlights the cluster containing highly similar protocadherin termini. **(D)** Binding scores mapped onto AlphaFold structure of *γ*-protocadherin A3 (UniProt accession: Q9Y5H0). p-value shows the probability of observing a binding site of this strength or stronger. The color scale corresponds to E. **(E)** Heatmap showing Binding scores of binding sites in the termini of protocadherins from cluster 1 in C. **(F)** Result of recurring fixed-length pattern motif discovery, identifying motifs enriched in non-AH binding sites compared to non-binding regions using STREME (Bailey, 2021) (see methods). **(G)** Density plot showing the location of the identified motif within binding sites, relative to the center of the binding site. **(H)** Overrepresentation analysis for PANTHER classes associated with proteins containing the motif identified in G, compared to all TMPs on the array. Point sizes and numbers show log_2_ fold change. p-values corrected using the false discovery rate. **(I)** Heatmap showing Binding scores for the motif in G found in voltage-gated ion channels enriched in H. **(J)** Binding scores mapped onto AlphaFold structure of CACNG2/Stargazin (UniProt: Q9Y698). Generated with MemRIDRdb. p-value shows the probability of observing a binding site of this strength or stronger. Binding site in CACNG2/Stargazin aligned to the cell penetrating peptides R9, TAT and penetratin. **(K)** Representative images showing immuno-staining for FLAG-Tac (Tac), FLAG-Tac-Hecate (Tac-Hecate), FLAG-Tac-G-shuffle (Tac-G-shuffle) and FLAG-Tac-Stargazin (Tac-Stargazin) following 30 min of internalization in the presence of recycling inhibitor monensin. Magenta, internalized; Green, surface. Scale bar 20 µm. **(L)** Quantification of K normalized to Tac. Median ± 95% CI of three independent experiments of at least 19 cells, total N>98 cells per condition, Kruskal-Wallis test with Dunn’s correction for multiple comparisons. *p-value ≤ 0.05, ****p-value ≤ 0.0001. ns>0.05. See also Figure S16.

To this end, we screened all non-AH binding sequences for short significantly enriched motifs in binding sites versus the non-binding background. Interestingly, this revealed that 499 binding sites in 452 TMPs contain motifs characterized by long stretches of arginine and lysine residues found distributed around the center of binding sites (∼9 amino acid long, consensus RKKKKKKKK) (Figure 7F-G). Apart from being able to interact with negatively charged lipid headgroups, such sequences strikingly resemble cell-penetrating peptides (CPPs). CPPs are positively charged peptides, which can modulate membranes and gain entry to the cell interior^53^. Examples of such CPPs include TAT (GRKKRRQRRRPQ), R9 (RRRRRRRRR), and penetratin (RQIKIWFQNRRMKWKK). In addition to identifying the existence of cationic motifs in cadherins, overrepresentation analysis highlighted such motifs in voltage-gated ion channels (Figure 7H). Among the 27 identified voltage-gated ion channels housing cationic motifs (Figure 7I) we encountered stargazin (transmembrane AMPAR regulatory protein (TARP) *γ*2) (Figure 7J). The identified region corresponds to a prominent motif that modulates the interaction of stargazing with PSD95 by tethering the stargazin tail to the membrane. The interaction is regulated by phosphorylation, which controls synaptic clustering of AMPA receptors and thereby synaptic plasticity^6,54–56^. The known functional role of the cationic motif in stargazin suggests that these sites play a similar role in the other identified voltage-gated ion channels. More broadly, we find that cationic motifs are overrepresented in TMPs associated with changes in membrane potential, such as during synaptic transmission and muscle contraction, or in TMPs related to cell adhesion and outgrowth (Figure S16).

We have previously shown that membrane-binding AHs can drive constitutive internalization of TMPs^24^. To evaluate whether cationic motifs act in a similar fashion, we fused the stargazin binding sequence to Tac and measured the rate of internalization by antibody feeding. As a reference, we fused the highly charged (+9) AH Hecate to Tac, as well as the shuffled derivative of Hecate, G-shuffle, to isolate the effect of charge alone. As previously reported, Hecate increased Tac internalization roughly 10-fold (Figure 7K-L), while G-shuffle largely abolished this effect (Figure 7K-L). Interestingly, G-shuffle still significantly induced Tac internalization roughly 1.5-fold, an effect we also observed for stargazin (Figure 7K-L). This suggests that short patches of cationic amino acids change the dynamics of TMPs. However, cationic binding sequences alone are not potent drivers of internalization compared to AHs. As shown for stargazin (Hafner *et al.*, 2015), regulatory mechanisms that rely on membrane interactions of short cationic motifs may have implications for understanding TMP clustering.

## Discussion

In this study, we generated a proteome-wide map of membrane binding sites in IDRs of human TMPs. Using this map, we demonstrate that the majority of IDRs contain membrane binding sites, primarily driven by AHs and less frequently by cationic motifs. We highlight the existence of AHs in channels and transporters and draw parallels to the amphipathic helix 8 of GPCRs. Such multi-pass TMPs contain most of the identified membrane binding sites. Although 1TM TMPs are largely devoid of membrane binding sites, we identify AHs in the short cytoplasmic tails of glycosyltransferases and demonstrate that these confer their localization to the Golgi apparatus. AHs are also known to help stabilize membrane curvature in MSAPs, and here we confirmed such known AHs and identified new AHs in most members of the superfamily, including in mitochondrial and peroxisomal proteins. Interestingly, AHs were prominent among mitochondrial and peroxisomal TMPs, which have an evolutionarily distinct origin from the rest of the endolysosomal system, hinting at an ancient role for AHs in TMPs predating the eukaryotic cell. This is supported by our finding that AHs are under higher evolutionary constraint and are associated with more pathological SNPs than the remaining IDRs. Additionally, several pathological SNPs disrupt such AHs and give rise to a wide range of diseases. Investigating non-AH binding sites, we identified stretches of basic residues as major drivers of membrane binding. Regions with such properties have been shown to play regulatory roles in cadherins, stargazin, synaptotagmin-1 and VAMP2^6,8,52,57^. Taken together, we present a map of membrane binding in IDRs of TMPs that covers membrane binding regions of known function and vastly expands their number, enabling the characterization of new and unconventional mechanisms for regulation of TMP function.

Understanding IDRs constitutes a major challenge in biology due to their unconventional sequence–structure–function relationships^3^. Here, we describe a ubiquitous role for specific parts of IDRs in TMPs as membrane-binding entities. Although not attributed to direct interactions with membranes, IDRs have previously been shown to modulate membranes by crowding^58^. When IDRs are crowded on membrane surfaces, steric and electrostatic repulsion drives the membrane to bend away from the IDR, such that the area available per IDR is increased. This process can lead to the formation of protein-coated membrane buds and tubules, prevent TMP internalization, and even drive membrane fission^59–61^. It would be valuable to explore an interplay between crowding and membrane binding of IDRs, where coupling of sites within long IDRs to the membrane could increase the crowdedness in proximity to the membrane, amplifying the curvature sensitivity of IDRs.

By bioinformatic analysis, we show that membrane binding in IDRs of TMPs can generally be divided into positively charged motifs and AHs. Characterizing the identified AHs, we find that aromatic residues play a central role in membrane binding and are often located centrally in the hydrophobic face. Biochemically, this is distinctly different from the positions in transmembrane helices (even when amphipathic in nature), where aromatic residues cap the helices to position themselves in the lipid–water interface^62^. Furthermore, we find that charged amino acids and their distribution within the AH also play a major role in the interaction with the lipid bilayer. Generally, we find that positive charges favor the interaction of AHs with membranes, whereas negative charges oppose the interaction as anticipated. This is in line with recent findings that interaction with negatively charged amino acids induces a large dehydration of membrane bilayers, while cationic amino acids arginine and lysine fluidize the membrane^63^. Recent molecular dynamics (MD) simulations find terminal charges to be the initial point of contact for AHs through an electrostatic interaction^64^. Whereas, the folding of AHs could be an iterative process, with folding and merging of lipid defects happening gradually^65,66^.

We find that AHs in TMPs are under significant evolutionary constraint, and that missense mutations in AHs more frequently result in disease, compared to the remaining IDRs. Several such mutations result in the disruption of AHs by, e.g. inserting charged residues in the hydrophobic face or vice versa. Examples of such substitutions can be found in channels and transporters, including L11P in ABCA4, W57G in CFTR, and 5 different substitutions of R553 of KCNQ2. The ABCA4 L11P mutation, identified in patients with Stargardt disease, prevents cell surface expression and thereby ABCA4 dependent ApoA1 cholesterol efflux^67^. Similarly, the rare CFTR W57G mutation results in minimal mature CFTR on the cell surface and responds poorly to current corrector drugs^68^. Substitutions of R553 in KCNQ2 underlie the development of Epileptic Encephalopathy and have been suggested to disrupt binding of the membrane lipid phosphatidylinositol 4,5-bisphosphate, which in turn prevents current potentiation^69^. Furthermore, mutations in this position reduce channel enrichment on the axonal surface, suggesting that trafficking is also impaired^69^. The AH could account for this dual role in KCNQ2, by driving efficient forward trafficking of the channel, as well as coordinating PIP2; as seen for the amphipathic helix 8 of the angiotensin receptor^24,70^. Substitutions of R553 also disrupt calmodulin binding, and it should be noted that the AH might also act as a multimerization domain^71^. Collectively, these studies suggest that membrane-binding AHs may play a critical role in proper TMP trafficking and maintenance of physiological function, and that mutations in these AHs underlie a spectrum of inherited pathologies.

Given the distinctive properties of the identified AHs in TMPs and the diverse membrane landscapes they encounter, they could serve multiple roles, including anchoring of terminal tails, membrane curvature induction, stabilization, sensing, or membrane fission. For highly curved membrane structures, such as synaptic vesicles, insertion of AHs could act to lower the membrane energy by relieving strain on a membrane that is far from its preferred curvature of zero^72^. For low-curvature membranes, such as the plasma membrane, AH insertion would have the inverse effect, destabilizing the membrane by introducing more strain, which could be the mechanism facilitating endocytosis^24^. Conversely, membrane binding regions within TMPs might also facilitate membrane fusion events. We identify a highly significant membrane binding site in the juxtamembrane domain of VAMP2, which has been reported to play a crucial role in the regulation of SNARE complex-mediated vesicle fusion. This region has been shown to act as a membrane-destabilizing peptide that is necessary for membrane fusion to occur^8^. Remarkably, the CPP TAT can replace the juxtamembrane domain of VAMP2 and rescue vesicle fusion^8^, highlighting the functional importance of the CPP-like nature of the juxtamembrane domain. The requirement of such membrane-destabilizing CPP-like regions is likely a common feature of membrane-fusion-inducing proteins, and similar regions in other proteins can likely be identified in our proteome-wide screen.

Here, we reveal 499 motifs in 452 TMPs that appear to interact with membranes via stretches of cationic amino acids. Apart from VAMP2, such regions have been shown to act as functional sites in stargazin and synaptotagmin-1, where they act as a regulatory mechanism for protein-protein and protein-membrane interactions by tethering cytosolic termini to the membrane and regulating its extension into the cytoplasm^6,7^. Additionally, membrane binding by similar motifs has been identified in *γ*-protocadherins^52^. However, the role of this interaction in *γ*-protocadherins remains unclear. The large number of identified regions allowed us to derive a consensus binding motif for such interactions consisting of up to nine cationic residues. The striking resemblance of the motif to CPPs suggests that such sites might play more advanced roles in TMPs than acting solely as distance rulers.

In summary, we have created a proteome-wide map of functional membrane binding sites in IDRs of human TMPs and highlight the widespread existence of membrane-binding AHs as well as cationic motifs. This map not only will aid the discovery of membrane-interacting regions important for TMP function but also provides valuable data to improve our general understanding of IDRs. Finally, modification or manipulation of the mapped membrane interacting regions may serve as strong research tools or even as putative pharmaceutical leads.

### Limitations of the study

In an attempt to identify the widest range of binding sites, we probed the array with a highly heterogeneous lipid extract from bovine brain. This came with the drawback that the extract might not represent any specific organellar lipid composition. Additionally, the extract was high in phosphatidylserine and cholesterol, giving the liposomes a plasma membrane bias and a higher negative charge. Furthermore, the sequences probed for membrane interaction may not be sterically available in the context of the protein or represent regions involved in protein-protein interactions. Therefore, the results must be interpreted with caution and the physiological importance of the reported membrane binding regions should always be verified in an appropriate system. Also, the 16-mer length of the peptides limits the resolution of the setup and may miss interactions based on complicated longer binding modes. Finally, cytosolic IDRs were determined using UniProt.org and predictions; thus, false positive regions might exist in the dataset as well as missing IDRs.

## Acknowledgements

We thank Claus Schafer-Nielsen (Schafer-N ApS) and Paola Saporito for their guidance during the setup and probing of the arrays. We thank the Core Facility for Integrated Microscopy at the University of Copenhagen for help with the TEM imaging. This project was funded by Danish Research Counsil (DFF), Medical Sciences, FP 1 (2034-00395A) (to K.L.M.); Lundbeckfonden, Lundbeck Foundation Experiment (R324-2019-1806) (to K.L.M.); Lundbeck Foundation Post Doc grant (R347-2020-2339) (to A.H.L.).

## Author Contributions

**Joscha Rombach:** Conceptualization, Methodology, Software, Formal Analysis, Investigation, Data Curation, Writing – Original Draft, Writing – Review & Editing, Visualization. **Tommas Theiss Ehler Nielsen:** Conceptualization, Methodology, Software, Formal Analysis, Investigation, Data Curation, Writing – Review & Editing, Visualization. **Junior Agenant:** Software, Investigation. **Andreas Haahr Larsen:** Writing – Review & Editing. **Jan Hendrik Schmidt:** Investigation, Writing – Review & Editing. **Kenneth Lindegaard Madsen:** Conceptualization, Methodology, Investigation, Resources, Writing – Review & Editing, Visualization, Supervision, Project Administration, Funding Acquisition.

## Supplemental information

Document S1. Raw data file for array replicate 1

Document S2. Raw data file for array replicate 2

Document S3. SnapGene DNA file of pcDNA3 Flag-Tac

Table S2. Excel file containing additional data too large to fit in a PDF related to Figure 5A

Image S1. Full image of array replicate 1

Image S2. Full image of array replicate 2

## Supplementary table titles and legends

**Table S1.**
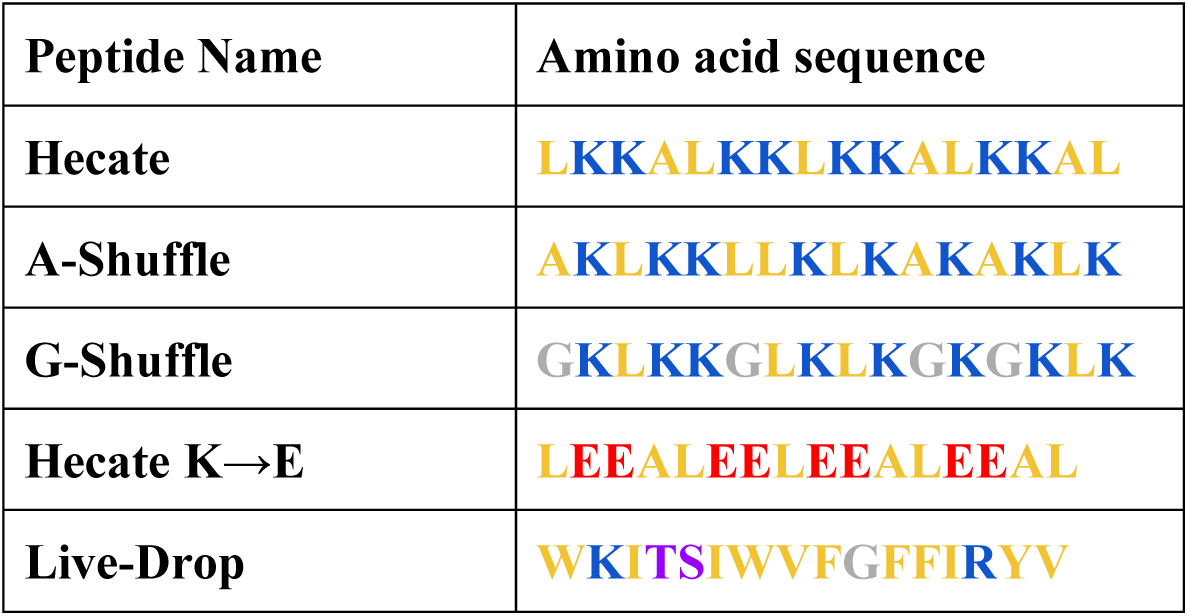
Control peptides on array. Amino acid color code: yellow; hydrophobic, blue; cationic, red; anionic, purple; polar, grey; glycine.

## Supplementary figure titles and legends

**Figure S1.**
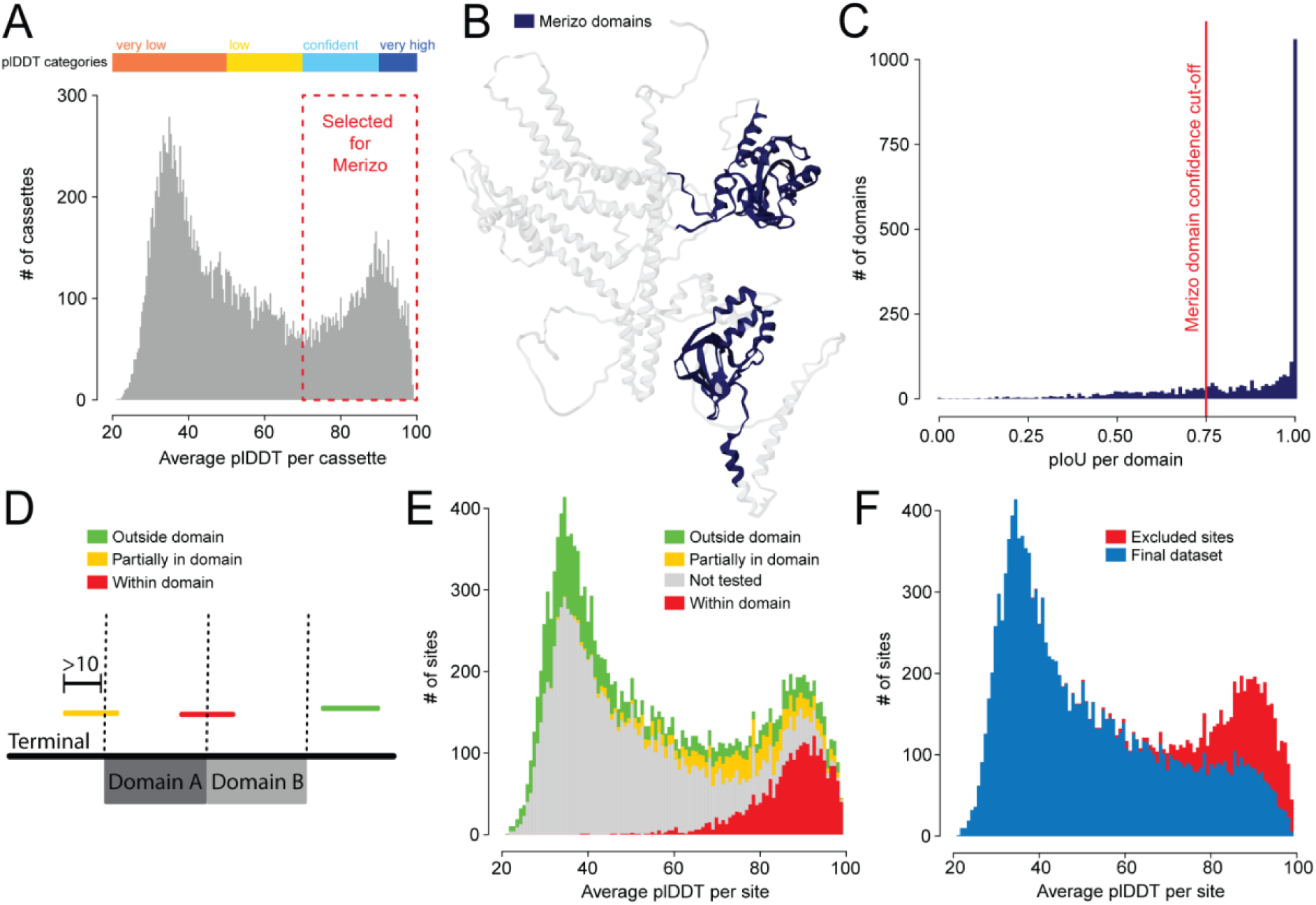
Domain Filtering using AlphaFold2 and Merizo. **A)** Histogram showing the distribution of average AlphaFold plDDT values of 40mer cassettes overlapping by 50% from termini pre-filtered for annotated domains. Color bar shows AlphaFold plDDT categories as set by Jumper et al.^73^. Red dashed box highlights cassettes selected for subsequent domain prediction using the Merizo model. (**B)** Example of domains predicted in the AlphaFold2 model of the Potassium voltage-gated channel subfamily H member 1 (AF-O95259-F2) by the Merizo model. Domains are highlighted in blue. **C)** Histogram showing the distribution of the intersection over union (pIoU) values from the Merizo model, an estimate for predictive confidence of the domains. Red line indicates the pIoU confidence threshold used to select true positive domains. **D)** Illustration showing how binding site-domain overlaps in termini were characterized. **E)** Stacked histogram showing the distribution of average plDDT scores for binding sites grouped by domain overlap as described in D. **F)** Stacked histogram of the distribution of average plDDT scores for binding sites showing excluded sites in red.

**Figure S2.**
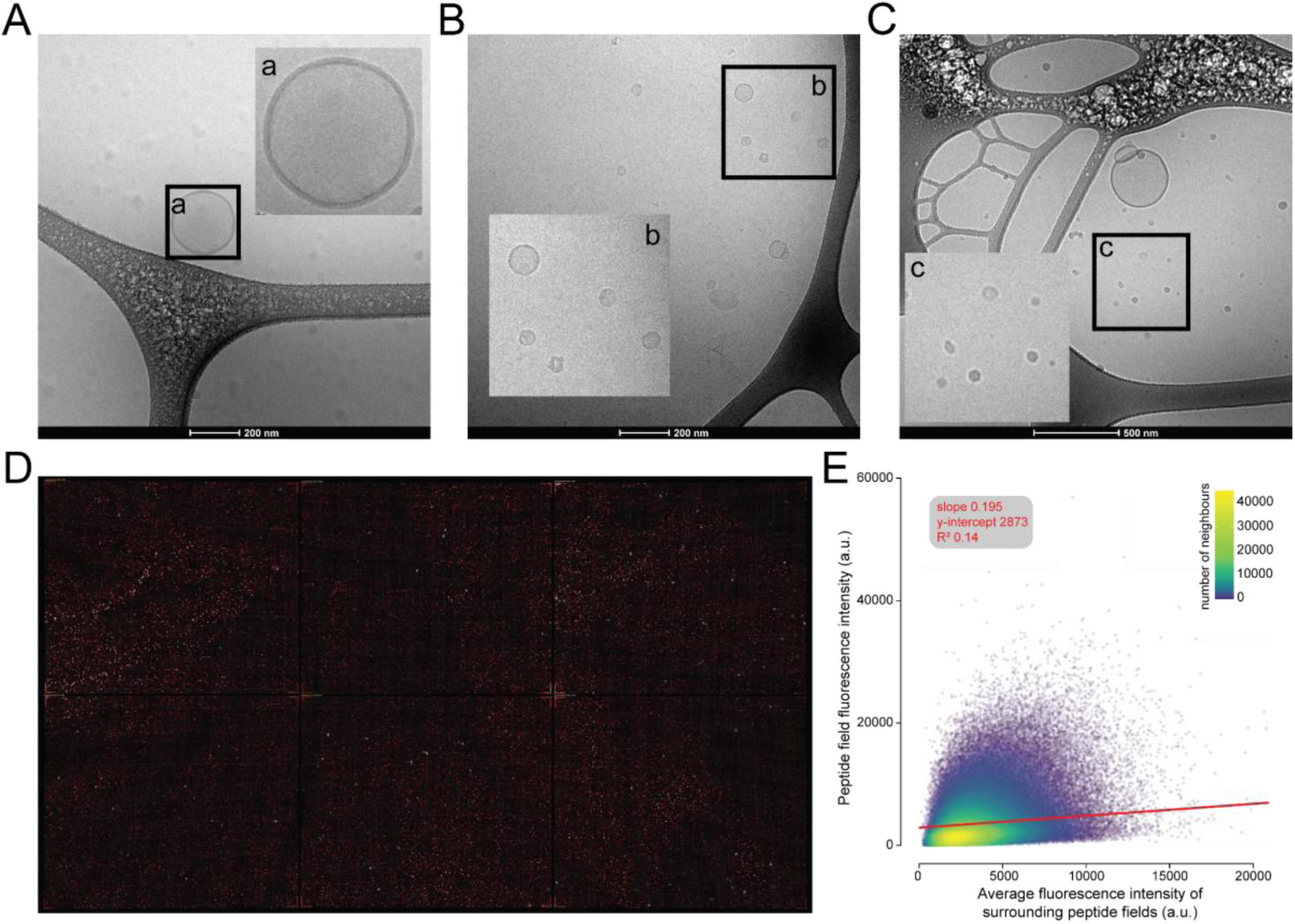
Cryo-TEM images and fluorescence of DiD-labeled liposomes. DiD liposomes were extruded to <1000 nm. **(A-C)** Representative Cryo-TEM images of DiD liposomes. **(A+B)** Scale bar 200 nm. **(C)** Scale bar 500 nm. **a-c** are inserts and the corresponding box shows the area zoomed in on. **D)** Raw 16-bit image of DiD-liposome fluorescence on array replicate 2 overlaid with a grid (gridline width = 10 *µ*m). **E)** Point plot showing how fluorescence intensity of surrounding fields affects field fluorescence. Color indicates the number of neighboring points with a smoothing bandwidth (i.e. within a radius) of 1000 a.u.. Red line shows the result of linear regression; slope: 0.195, y-intercept: 2873.46, R^2^: 0.14.

**Figure S3.**
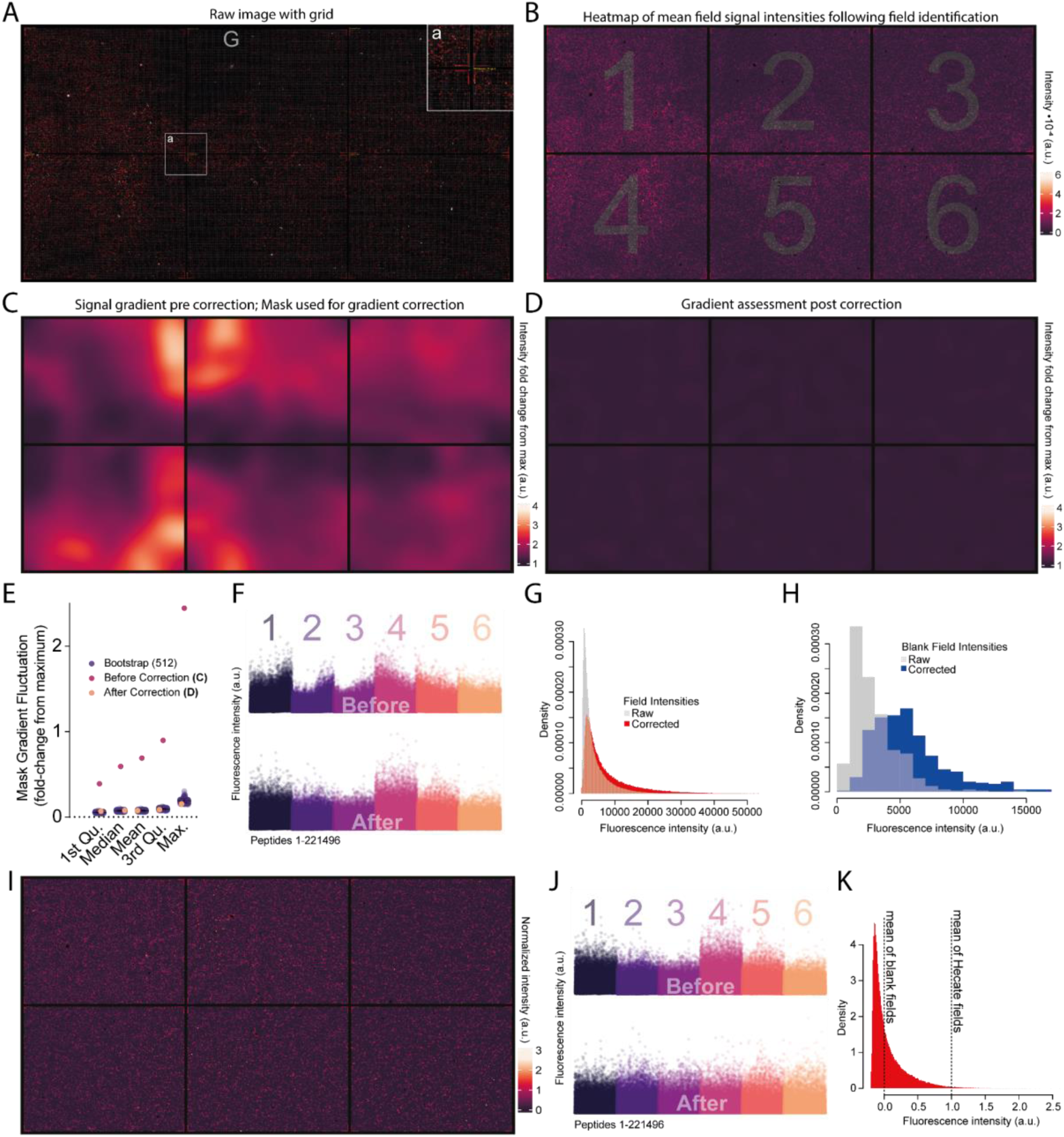
Gradient correction and signal normalization. **(A)** Raw 16-bit image of DiD-liposome fluorescence on array replicate 1 overlaid with a grid (gridline width = 10*µ*m). Insert **a** highlights 17 fields in each corner of all sectors containing a small subset of the positive control peptide; the strong amphipathic helix Hecate (amino acid sequence: LKKALKKLKKALKKAL). **(B)** Mean values of fluorescence intensities in each field plotted as a heatmap. Gray numbers indicate the six sectors. **(C)** Result of kernel smoothing using the Nadaraya/Watson normalization of the kernels used as a mask to correct for fluorescence gradients on the array arising due to washing and drying steps following incubation with DiD-liposomes. Intensities normalized to maximum. **(D)** Result of kernel smoothing following gradient correction, showing only expected minor mask gradient fluctuations. Data normalized to max. **(E)** Distribution of intensities following kernel smoothing before and after gradient correction, as well as the distribution of intensities from 512 uniform mock arrays with completely random distribution of intensities created by bootstrap sampling from the array in B. All kernel smoothing results are normalized to max. **(F)** Fluorescence intensities of individual peptide fields on the array before (top) and after (bottom) gradient correction. Colors indicate sectors. **(G)** Histogram showing the distribution of fluorescence intensities before and after gradient correction. **(H)** Histogram that shows the distribution of fluorescence intensities from empty fields on the array before and after gradient correction. **(I)** Heatmap showing the gradient-corrected fluorescence intensities on the array. **(J)** Fluorescence intensities of individual peptide fields on the array before corrections (top) and after gradient correction and sector normalization to Hecate and empty fields (bottom). Colors indicate sectors. **(K)** Histogram showing the distribution of fluorescence intensities after gradient correction and normalization.

**Figure S4.**
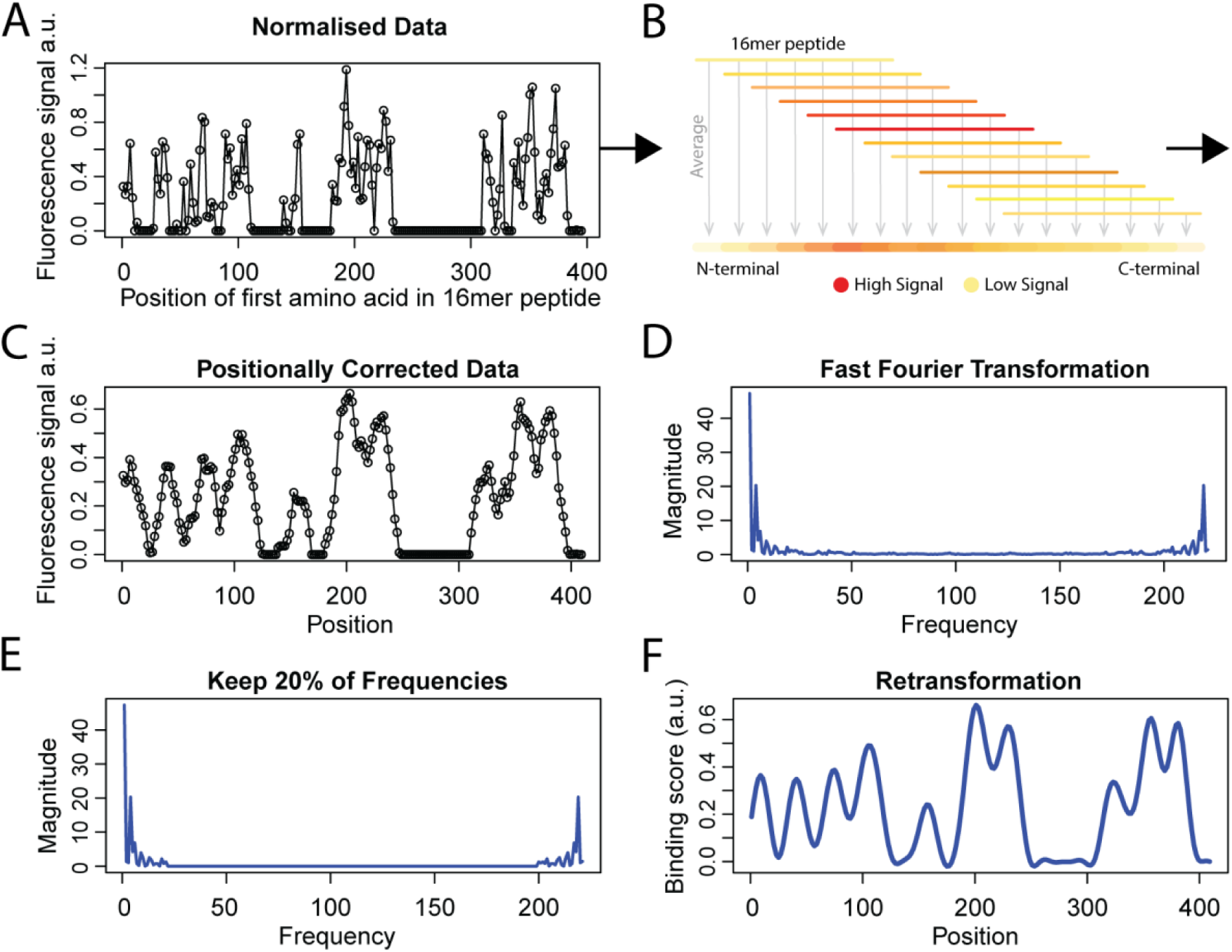
Calculating smoothed weighted-average fluorescence intensity scores (Binding-scores) **(A)** Example of terminal reconstruction from mean fluorescence intensities for individual peptide fields. Reconstruction of the cytosolic N-terminal of the Epidermal growth factor receptor-related protein, iRhom1, (UniProt: Q96CC6) from the gradient-corrected and normalized fluorescence intensities. Each point represents the first amino acid in each 16-mer peptide. **(B)** Schematic illustrating positional correction. Peptides are synthesized on the array with the C-terminal attached to the substrate on the glass slide. **(C)** Trace of reconstructed terminal following positional correction. **(D)** Fast Fourier transformation of trace in C. **(E)** Fast Fourier transformation after removal of 80% of frequencies. **(F)** Re-transformation of E resulting in the final trace. Binding score is therefore a normalized smoothed weighted-average fluorescence intensity.

**Figure S5.**
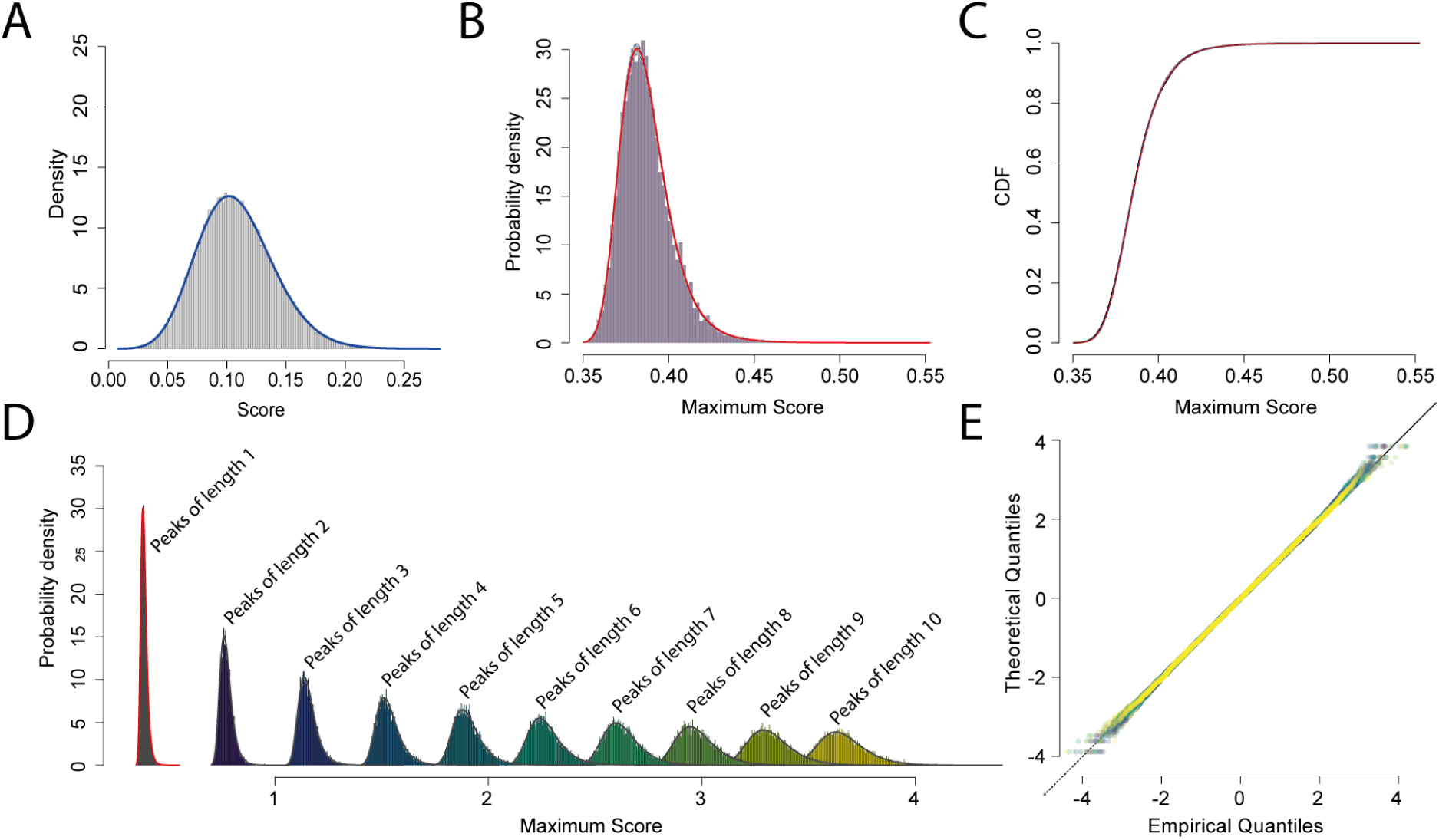
Fitting background distributions for statistical testing. **(A)** Mean binding scores of all terminals (histogram) fit with a power-normal curve (blue line). This distribution was used for imputation of missing binding scores removed due to artifacts (Figure 1C). **(B)** Example of a background distribution for peaks of length 1 for a terminal of length 1360 used for statistical testing. 10,000 random terminals of length 1360 were generated by bootstrap sampling fluorescence intensities of fields on the array. The fluorescence intensities were assembled into ‘mock’ terminals and binding scores calculated as done for actual IDRs (Figure S4). The maximum binding score for each ‘mock’ terminal was extracted. 100 sub-samplings of the resulting data were fit to power-normal distributions and density functions were calculated. Red line indicates 50th quantile, blue dashed lines indicate 5th and 95th quantiles and gray lines are all fits. Such power-normal probability curves were used to calculate the significance of binding scores in actual IDRs. **(C)** Cumulative density function of the 10,000 samplings described in B (black line). 100 sub-samplings of the resulting data were fit with a power-normal curve and the distribution function was calculated. Red line indicates 50th quantile, blue dashed lines indicate 5th and 95th quantiles, gray lines are all fits. **(D)** Example histograms of maximum binding scores as calculated in B for binding peaks of lengths 1 to 10. Data were fit with power-normal probability curves (red (same as in B) and black lines), which were later used to calculate the significance of peaks of the given lengths (Example Figure S6). **(E)** qq-plot for all fits as in D, but for lengths 1 to 200. Points colored using the viridis color scale by fit, from purple (length 1) to yellow (length 200).

**Figure S6.**
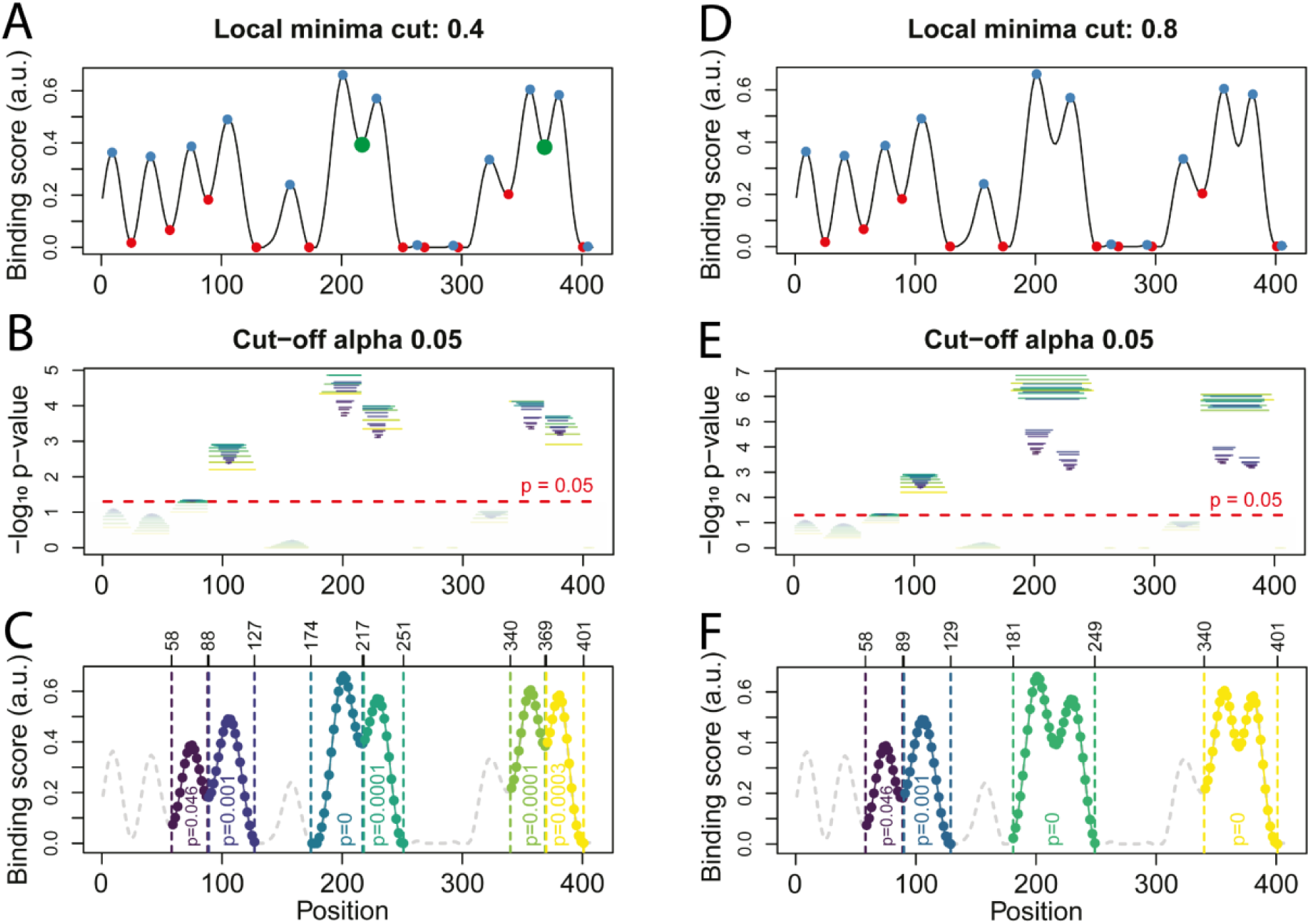
Peak identification and statistical testing. **(A)** Peak identification with a minima cut-off of 0.4; a peak is split if a local minima is <60% of the previous local maxima. red points; minima, blue points; maxima, large green points; vary depending on minima split threshold. **(B)** Example of statistical testing of each peak in a terminal with a significance cut-off of p-value = 0.05. Line length indicates the region tested (colors show site length, see Figure S5D), red dashed line; −log_10_(0.05). **(C)** Identified binding sites with corresponding p-values, colored by site. **D-F** Same as A-C, but with a local minima cut-off of 0.8. For this paper the local minima cut-off was set to 0.5 and significance threshold to 0.05 as default.

**Figure S7.**
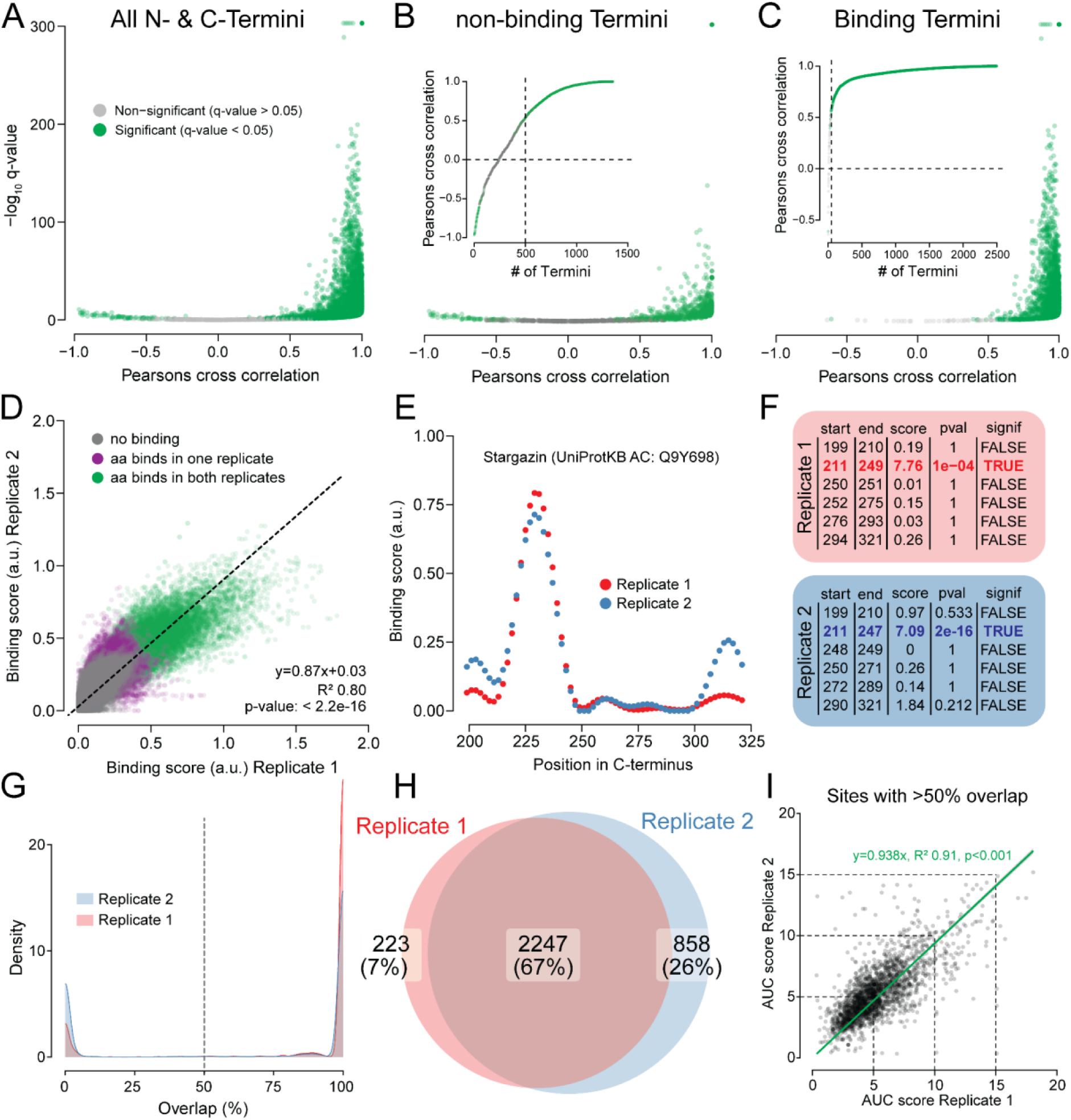
Liposome binding on the peptide array is robust and reproducible. **(A)** Results of Pearson’s cross correlation between corresponding IDR traces from replicate 1 and 2. p-values corrected for multiple testing with false discovery rate (q-value) **(B)** Results of Pearson’s cross correlation for IDRs in the non-binding group (p-value > 0.95). Insert shows results ordered by Pearsons cross correlation. **(C)** Results of Pearson’s cross correlation for IDRs that exhibit statistically significant membrane binding (p-value < 0.05). Insert shows results ordered by Pearsons cross correlation. **(D)** Correlation between binding scores for corresponding amino acid pairs in replicate 1 and 2. Dashed line is a linear fit; slope = 0.87, y-intercept = 0.03, R^2^ 0.80, p-value = 0. **(E)** Example of a corresponding binding trace from replicate 1 and 2. **(F)** Scores and p-values for each peak in the traces from E. **(G)** Density plot showing the degree of overlap between binding sites identified in replicate 1 and 2, and vice versa. **(H)** Venn diagram showing number of binding sites with an overlap >50% between replicate 1 and 2. **(I)** Correlation between AUC scores for sites with significant binding and more than 50% overlap between replicate 1 and 2. Green line is a linear fit, y=0.938x, R^2^ 0.91, p-value < 0.001.

**Figure S8.**
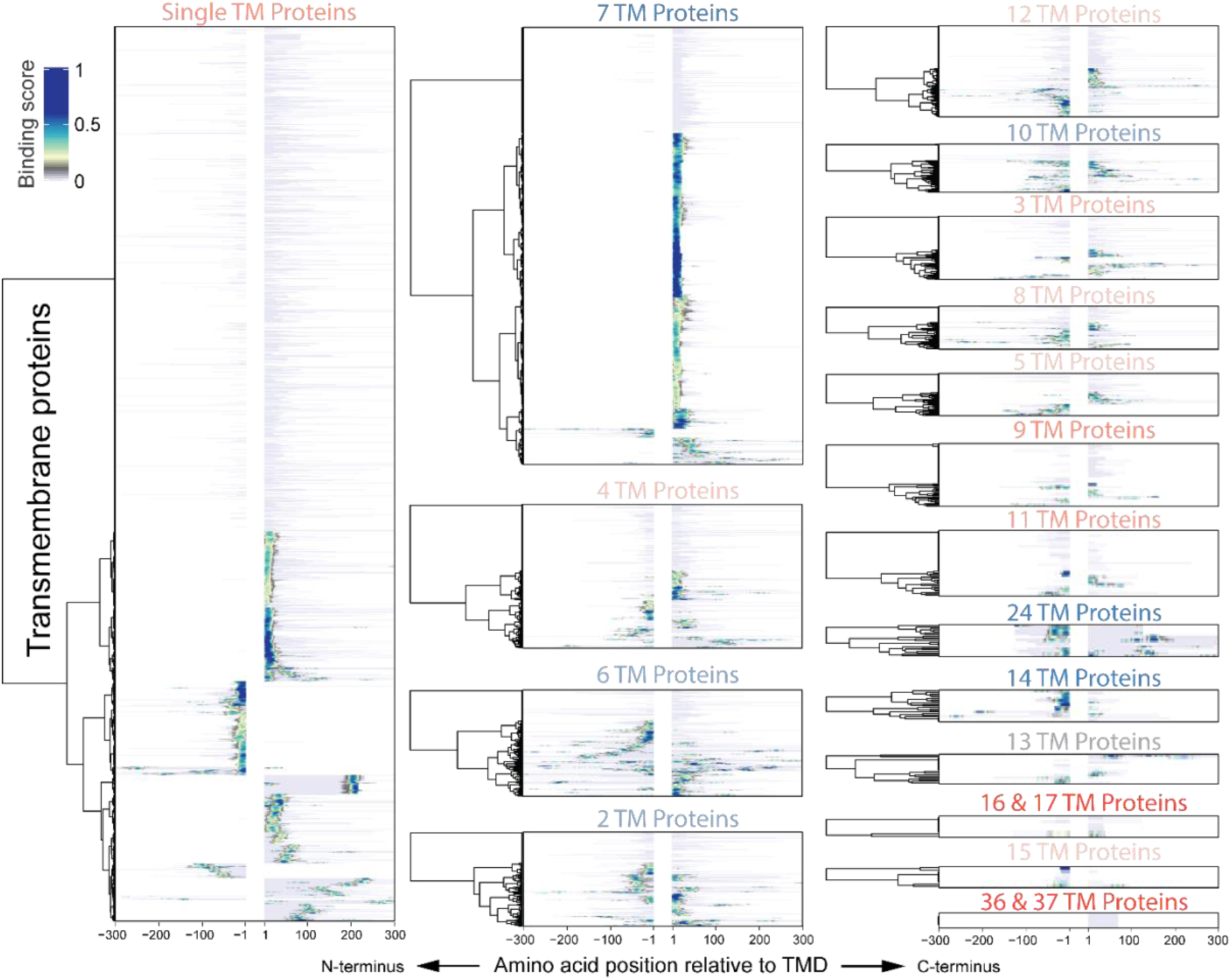
Membrane binding sites are a global attribute in IDRs from TMPs with diverse architectures. Heatmaps showing membrane-binding scores for binding sites in all TMPs, grouped by topology. Red and blue colors refer to the fold change from Figure 2G.

**Figure S9.**
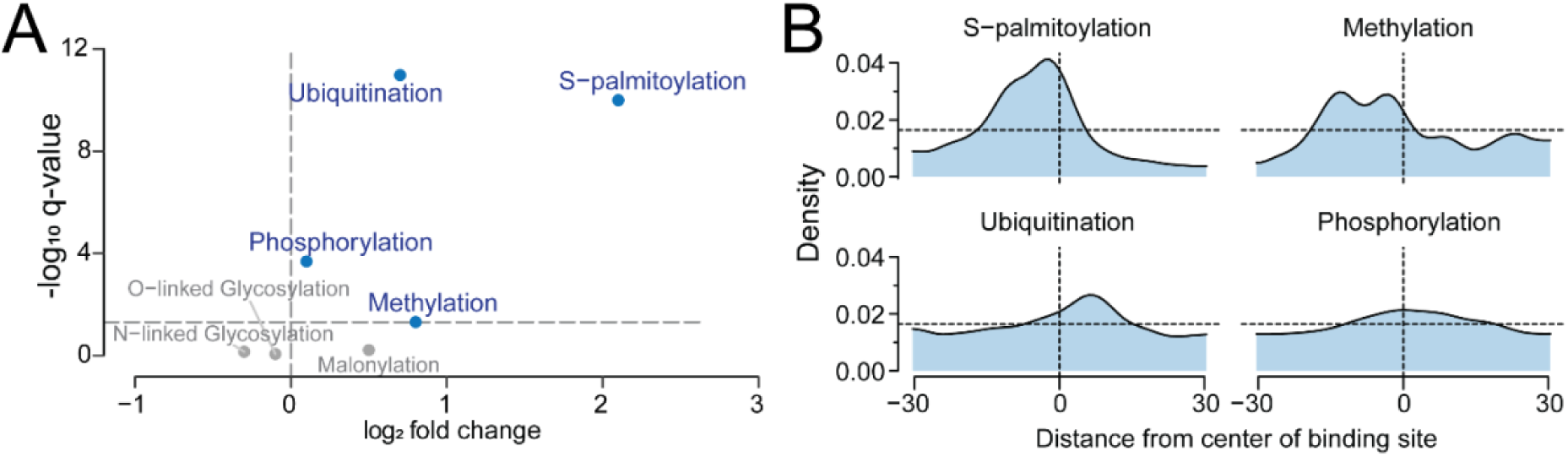
Post-translational modifications in binding sites. **(A)** Volcano plot shows the enrichment of post translational modifications contained within binding sites - non-binding sites. Fishers’ exact test, p-values corrected for multiple testing using the false discovery rate (q-values). **(B)** Probability density plots (n = 61, bandwidth = 3) showing the relative position probability of post translational modifications with respect to the center of binding sites extending also outside binding site proper. Vertical dashed lines indicate binding site centers. Horizontal dashed lines show the probability density expected for perfectly evenly distributed data. Data from dbPTM^74^.

**Figure S10.**
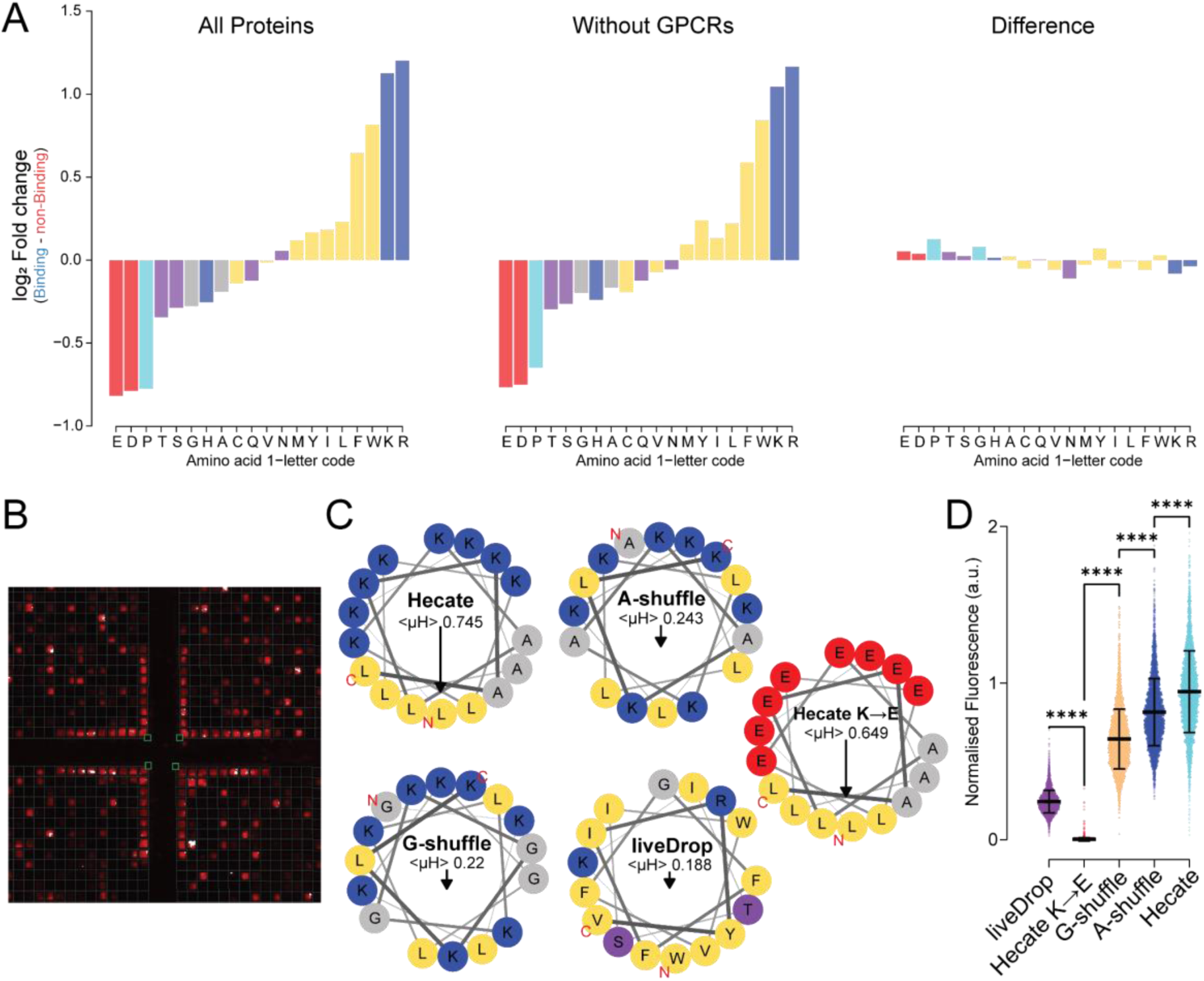
Cationic and hydrophobic amino acids drive liposome binding. **(A)** Amino acid enrichment in binding vs non-binding sites (left) Amino acid enrichment in binding vs non-binding sites excluding GPCRs (middle). Difference between Amino acid enrichment in binding vs non-binding sites including and excluding GPCRs (right). **(B)** Image of array highlighting the DiD-liposome binding to the 17 positive control fields containing Hecate synthesized in each corner of each sector on the array. **(C)** Helical-wheel representations of control sequences; Hecate, A-shuffle, G-shuffle, Hecate K→E and liveDrop. <µH>, mean hydrophobic moment. **(D)** Gradient corrected and normalized fluorescence intensity of peptide fields containing controls in E. Mean ± SD, One-way ANOVA with Dunn’s correction for multiple comparisons, ****p-value<0.0001.

**Figure S11.**
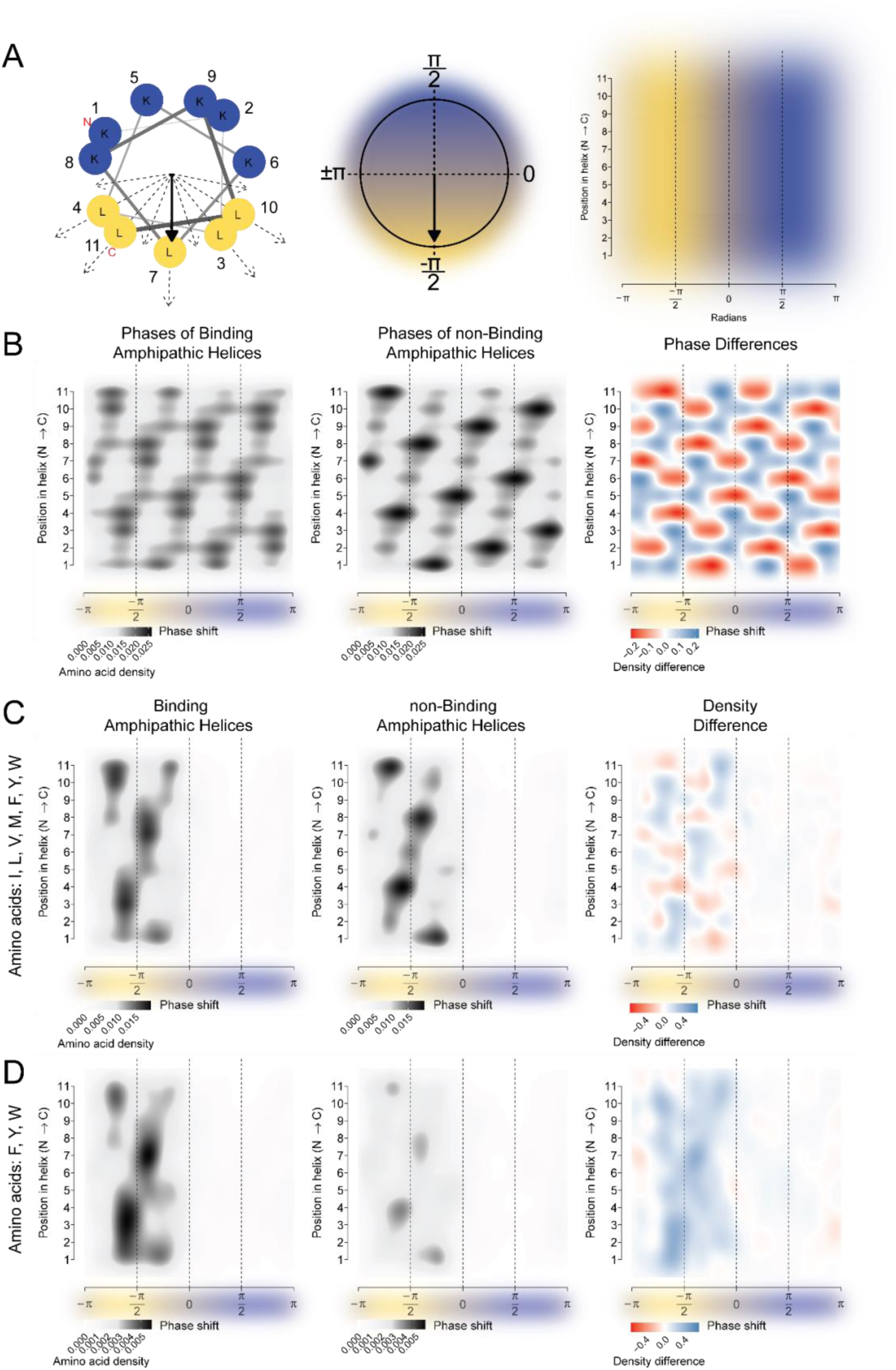
Amino acid distribution in liposome binding and non-binding amphipathic helices. **(A)** Schematic illustrating generation of 2D helical density plots. left: Helical wheel projection of an AH. Numbers indicate amino acid index from N-to C-terminus, dashed arrows show vectors representing the hydrophobicity of each amino acid, filled arrow is the mean hydrophobic moment vector. middle: Unit circle showing radians used to unfold helical-wheel projection into 1 dimension. right: Plot illustrating the final projection of the AH with radians on the x-axis and amino acid position in the helix on the y-axis. **(B)** Density plot showing the position of amino acids in AHs. *left*: Position of amino acids in liposome binding AHs (11mer with the highest mean hydrophobic moment from each site). *middle*: Same as left but for AHs that do not bind liposomes on the array. *right*: Relative density difference between amino acid positions in binding and non-binding AHs, log_2_(binding ⋅ non-binding^-1^). **(C)** Same as in B, but showing differences in density of amino acids I, L, V, C, M, F, Y, W. Amino acids weighted by their hydrophobicity according to their octanol/water partition^75^. **(D)** Same as C, but for amino acids F, Y, W.

**Figure S12.**
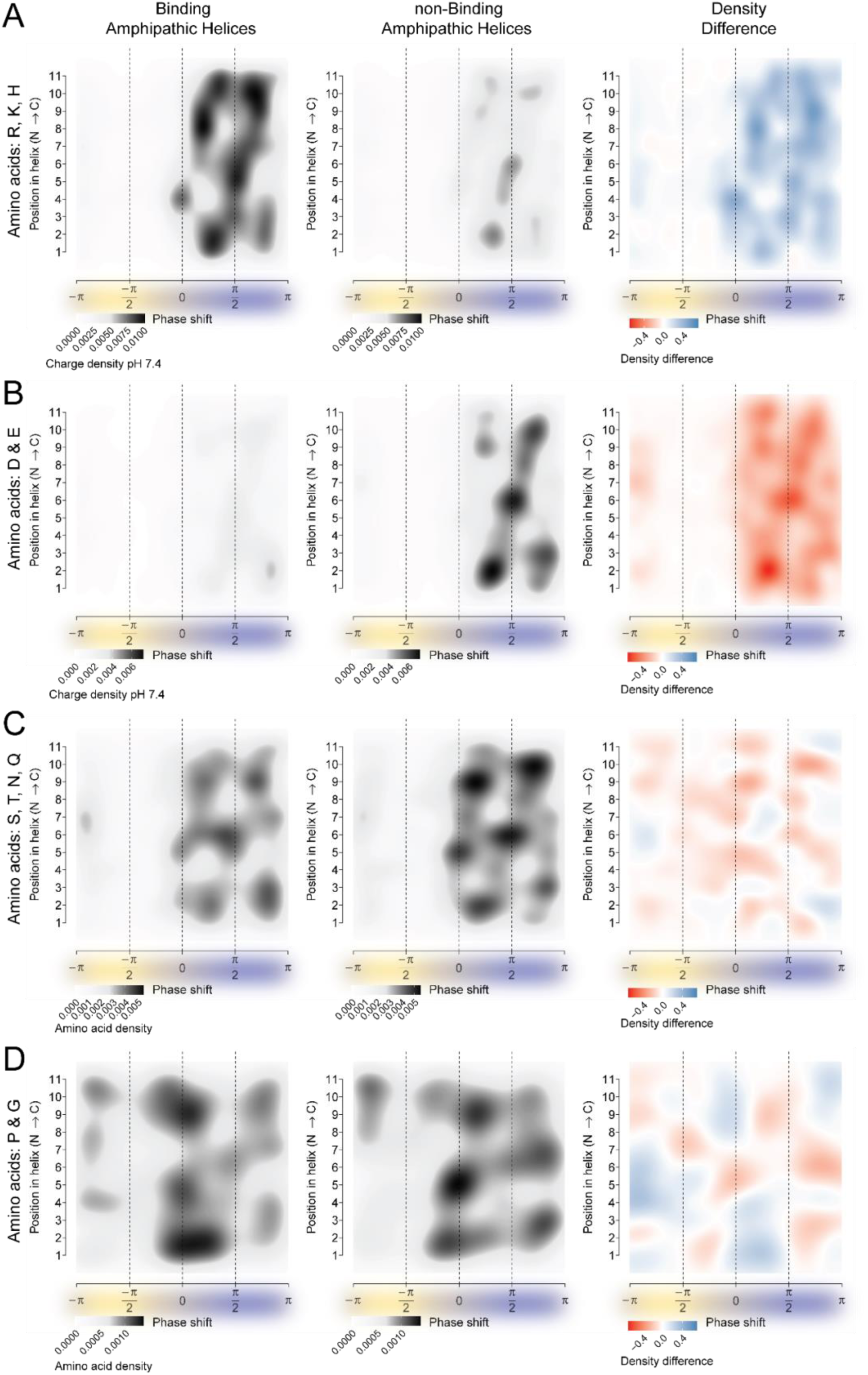
Amino acid distribution in liposome binding and non-binding amphipathic helices. **(A)** Density plot showing the density of charged amino acids R, K and H in AHs calculated at pH 7.4. *left*: Density in liposome binding AHs (11mer with the highest mean hydrophobic moment from each site). *middle*: Same as left, but for AHs that do not bind liposomes on the array. *right*: Density difference in binding and non-binding AHs, log_2_(binding⋅ non-binding^-1^). **(B)** Same as in A, but for amino acids D and E. **(C)** Same as A, but showing amino acid density for polar amino acids S, T, N and Q. **(D)** Same as C, but for amino acids with low helical propensity, P and G.

**Figure S13.**
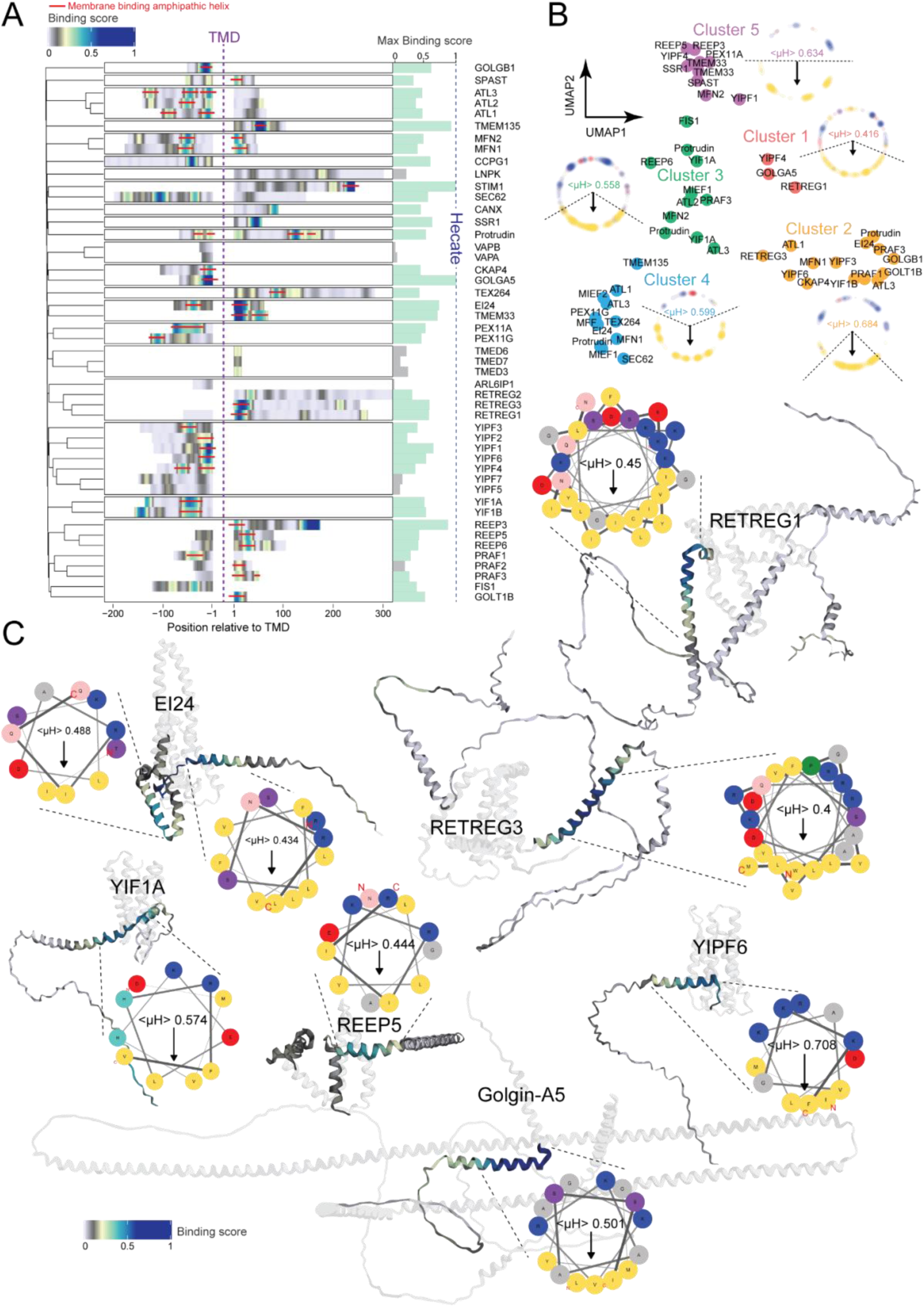
Amphipathic helices in membrane shaping adaptor proteins. **(A)** Heatmap showing binding scores in the termini of membrane shaping adaptor proteins (Angelotti, 2022). Membrane binding AHs highlighted with red lines. **(B)** UMAP showing the k-means clustering result of helical wheel-based hydrophobicity profiles for all AHs. Helical wheel density plots show the characteristics of AHs in each cluster. **(C)** Binding scores AHs mapped onto AlphaFold structures of selected MSAPs, including MSAPs with known AHs; REEP5 and RETREG1. Generated with MemRIDRdb. AHs are highlighted as helical wheel projections; <µH>, mean hydrophobic moment. See also Figure S14-15.

**Figure S14.**
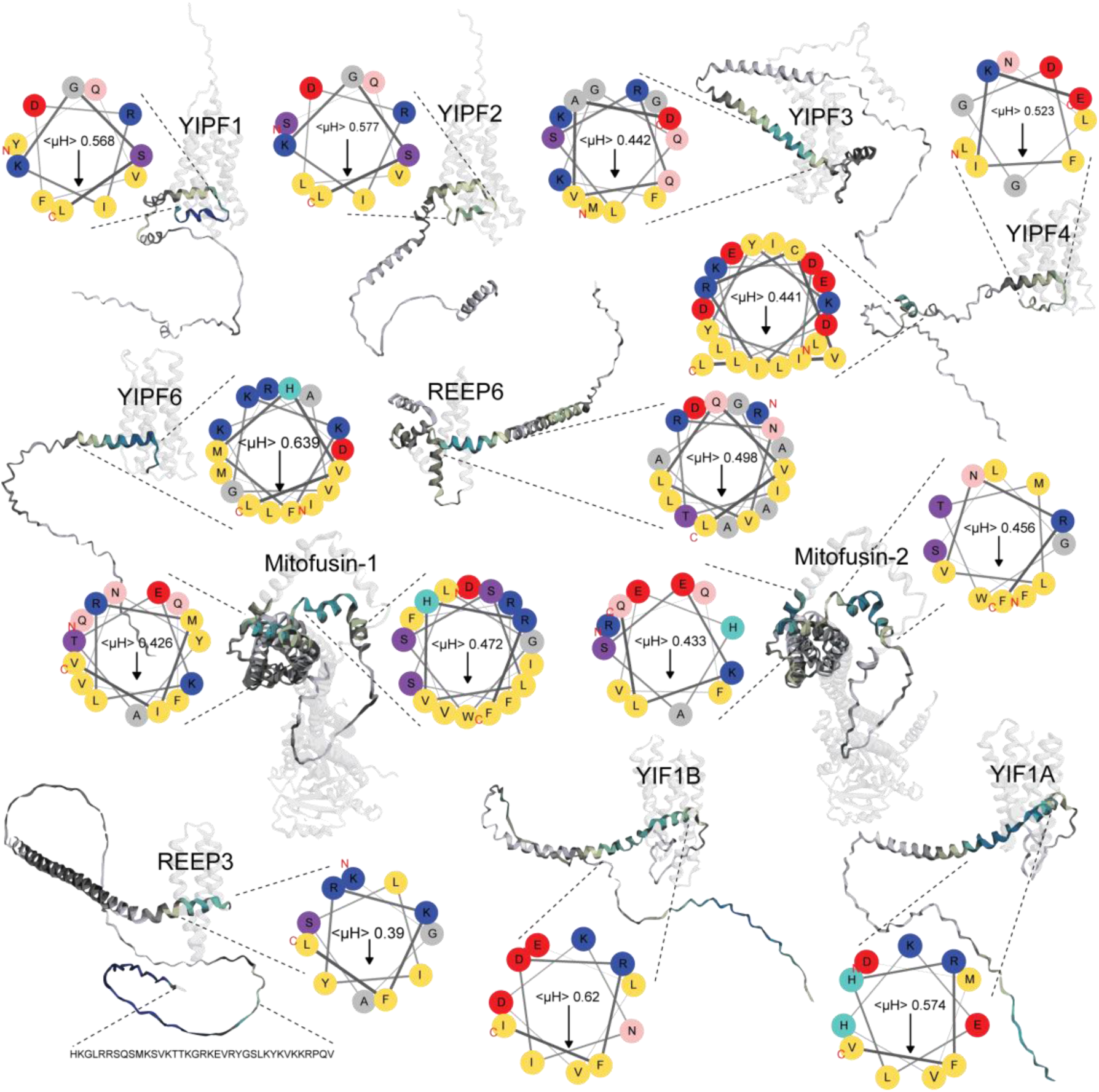
Examples of amphipathic helices in membrane shaping adaptor proteins. Binding scores mapped onto AlphaFold structures of membrane shaping adaptor proteins. AHs are highlighted and shown as helical wheel projections, including the calculated mean hydrophobic moment (<µH>). A binding site in REEP3 not predicted to be an AH is shown as sequence, bottom left. Generated with MemRIDRdb.

**Figure S15.**
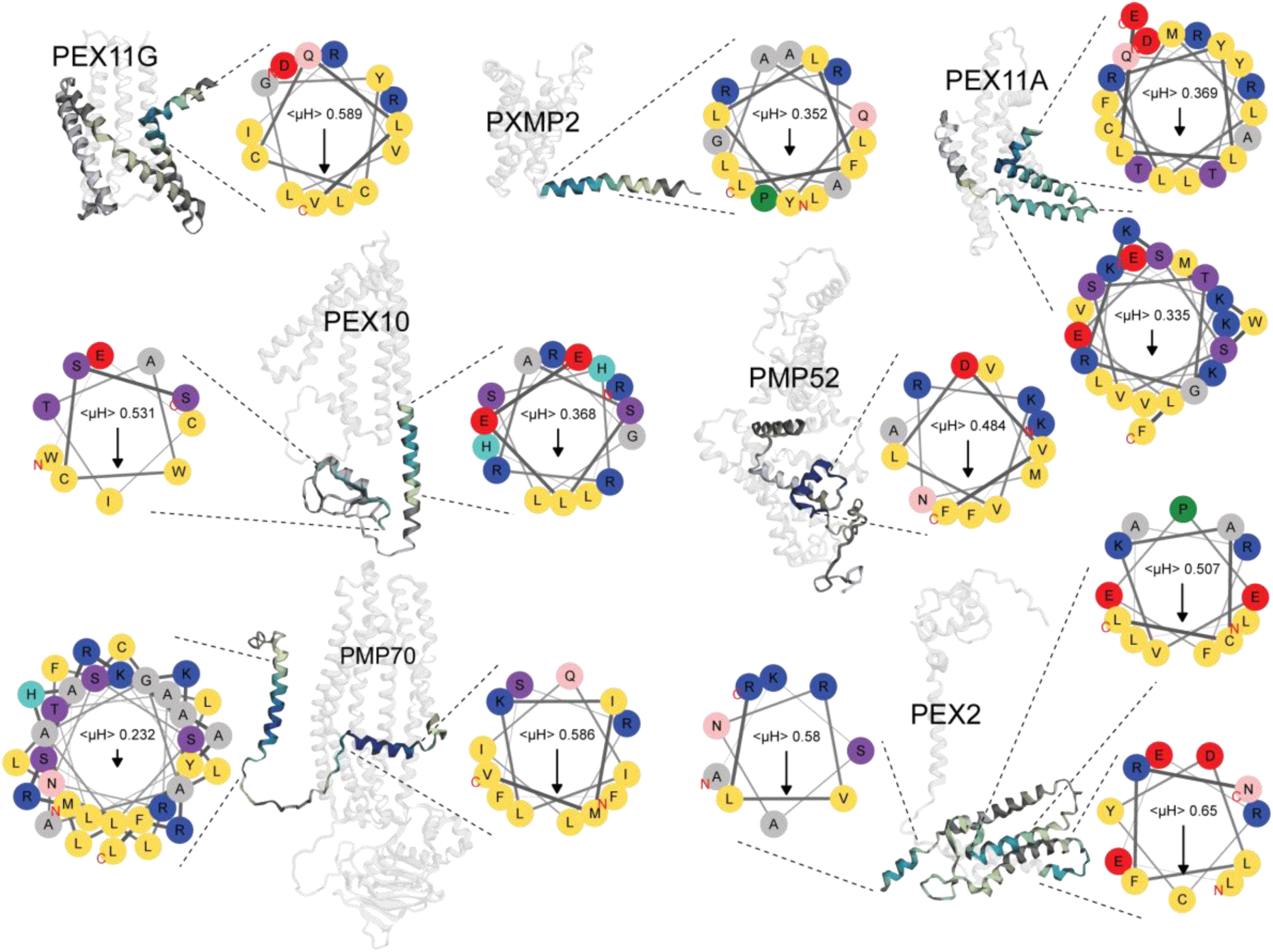
Amphipathic helices in Peroxisomal proteins. Binding scores mapped onto AlphaFold structures of peroxisomal transmembrane proteins. AHs are highlighted and shown as helical wheel projections including the calculated mean hydrophobic moment (<µH>). Generated with MemRIDRdb.

**Figure S16.**
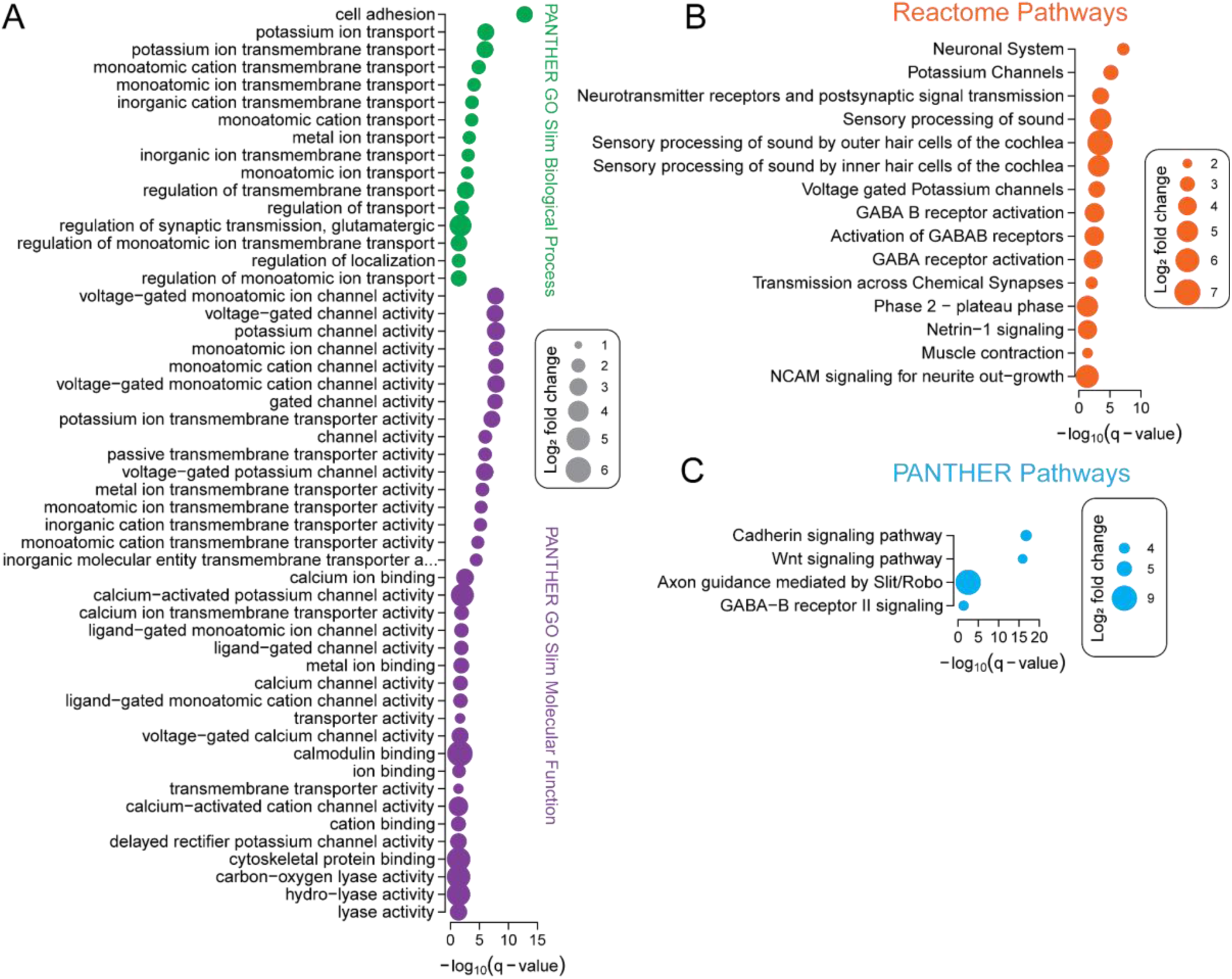
Cationic motifs are found in transmembrane proteins associated with changes in membrane potential and cell adhesion and outgrowth. **(A)** Overrepresentation analysis for PANTHER GO Slim molecular function and biological process terms associated with proteins containing the membrane binding cationic motif (consensus RKKKKKKKK), compared to all TMPs on the array. Point size shows log_2_ fold change, p-values corrected using the false discovery rate. **(B)** Same as in A, but for Reactome Pathway terms. **(C)** Same as in A, but for PANTHER Pathways.

## STAR Methods

### RESOURCE AVAILABILITY

#### Lead contact

Further information and requests for resources and reagents should be directed to and will be fulfilled by the lead contact, Kenneth Lindegaard Madsen.

#### Materials availability

Plasmids used in this study will be shared by the lead contact upon request, subject to the completion of a materials transfer agreement.

#### Data and code availability

- Images acquired of the arrays and the raw files generated from them have been deposited at GitHub. The link to the GitHub page is listed in the key resources table. Microscopy data (including selected regions of interest) in this paper will be shared by the lead contact upon request.
- All original code has been deposited at GitHub and are publicly available as of the date of publication. The link to the GitHub page is listed in the key resources table.
- Any additional information required to reanalyze the data reported in this paper is available from the lead contact upon request.

### EXPERIMENTAL MODEL AND STUDY PARTICIPANT DETAILS

#### Cell culture and cell lines

Human Embryonic Cells (HEK293) were cultivated in growth media (Dulbecco’s Modified Eagle Medium 1965 (DMEM) supplemented with 10% Fetal Bovine Serum (FBS) and 1% Penicillin/Streptomycin) and maintained at 37*^°^*C and 5% CO_2_ levels in humidified incubators. Cells were subcultured every 3-4 days (maximum passage 30) and never allowed to reach confluency above 95%. Cells were handled in sterile LaboGene ScanLaf Mars Pro safety cabinets and counted using a Countess automated cell counter (Invitrogen, C10281). Dead cells were identified using trypan blue. For routine passage or seeding, cells were detached from plates using 0.05% Trypsin-EDTA. Regarding sex, the HEK cells are female.

### METHOD DETAILS

#### Retrieval of Membrane Proximal N-& C-terminal IDRs

We extracted the complete set of Homo sapiens (taxid: 9606) gene products from the UniProtKB Swiss-Prot database with the Gene Ontology (GO) annotation “part_of integral component of membrane” (GO:0016021) from the EBI GO Annotation database (GOA) (https://www.ebi.ac.uk/GOA, 2020-08-17). This resulted in a dataset containing 6799 annotations to 5280 distinct gene products, which we exported as a Gene Product Association Data file. The UniProtKB accession numbers were used to extract their respective topological data from the Universal Protein Resource Knowledge-base (UniProtKB) (https://www.uniprot.org/). The following extraction, sorting and analysis of data was performed using the programming language R (v4.0.2). Gene products containing transmembrane and cytosolic N- or C-terminal domains were extracted from the dataset and N-terminal signal peptides and initiator methionines were removed. 1412 proteins were missing annotations concerning N- and C-terminal subcellular localization, which was predicted using TMHMM Server v. 2.0 (http://www.cbs.dtu.dk/services/TMHMM/)^76^. This resulted in 1167 successful predictions. Domains and intermembrane regions annotated in UniProt (Domain [FT], downloaded 08/03/2020) were removed. Sequences were additionally analyzed for domains (see section on domain filtering). The identified regions were removed and only the regions residing in proximity to transmembrane domains were kept. The resulting sequences were split into 16-mer cassettes overlapping by 14 amino acids (2 amino acid resolution), which resulted in 212,761 unique peptides.

#### Domain filtering

The termini of all proteins in the dataset were divided into cassettes of 40 amino acids overlapping by 50%. The cassette at the end of each terminal was always set to length 40. Terminals shorter than 40 amino acids were treated as one cassette. The average per-residue model confidence score (plDDT) of the AlphaFold structure was calculated per cassette (Figure S1A). A plDDT cut-off of 70 was used to select cassettes (Figure S1A). For the proteins with at least one cassette with an average plDDT > 70, domains were predicted with the Merizo model^77^ (Figure S1B). The Merizo model was run in Python (v3.11.5). An estimate for predictive confidence of the domains is the intersection over union (pIoU). Domains with a pIoU of 0.75 and a plDDT of 70 or higher were considered true positives (Figure S1C). Domains that fell within UniProt annotated termini were selected for subsequent filtering. For each termini the proportion within a domain was calculated. The sites that completely fell within a domain were filtered out and sites that completely fell outside a domain were included. Upon manual inspection of the results, domains that partially fell inside of the termini were most often transmembrane domains extending into the terminal or domains spanning over an α-helix. These domains were not used for site filtering. The termini that partially fell within a domain were flagged and included if they had at least 10 amino acids that did not fall inside of domains (Figure S1D-F). The analysis was performed in R (v4.3.2). Plots were generated using R packages **cowplot** (v1.1.3) and **ggplot2** (v3.5.0). Data was handled using **dplyr** (v1.1.4), **stringr** (v1.5.1), and **tidyr** (v1.3.1).

#### Generation of liposomes

Liposomes were prepared from bovine brain extract (Type I, Folch fraction I, Sigma-Aldrich) by following a standard hydration/extrusion procedure. The lipids were dissolved in chloroform. 2% DiD (w/w) was added for array experiments. Lipids were dried while rotating using argon-gas, creating a thin lipid film, followed by vacuum dehydration for at least 12 hours to remove residual chloroform. Lipids were rehydrated in sterile 200 mM D-Sorbitol solution (pH 7.4) to a final concentration of 0.5 mg/mL for array experiments. For cryo-TEM imaging lipids were rehydrated in sterile PBS (pH 7.4) to a final concentration of 1.25-2.5 mg/mL. The rehydrated lipids were put through a minimum of 8 “freeze-thaw” cycles; frozen using liquid nitrogen and thawed in a 40-50°C water bath. The liposomes were extruded through a 19 mm polycarbonate Whatman^TM^ Nucleopore^TM^ Track-Etched membrane with a pore size of 1 µm using a LiposoFast liposome extruder (Avestin). The liposomes were extruded at least 17 times. For the array incubations liposomes were diluted 1:100 in PBS (pH 7.4). For cryo-TEM imaging liposomes were diluted 1:10 in PBS (pH 7.4).

#### Cryo-TEM imaging

Lacey grids (Lacey F/SiO 300 mesh Cu) were glow discharged for 1 min with a CTA 010 device (Balzers Union). Samples were vortexed and 5 µL were added to the Lacey grids. Sample vitrification was done using a Thermo Scientific Vitrobot^TM^ Mark IV system (blot force: 2, blot time: 3 s, blot total: 1, wait time: 0 s, drain time: 0 s, humidity: 100%, temperature: 4*^°^*C) and plunge frozen in liquid ethane (cooled by liquid nitrogen). The grids were examined with a FEI Tecnai G2 20 TWIN at a voltage of 200 kV (grids were loaded and allowed to stabilize for 15 min prior to examination). Images were acquired with a bottom-mounted FEI High-Sensitive 4k x 4k Eagle camera.

#### Array synthesis, probing and scanning

Synthesis was performed by Schafer-N ApS, Lersö Parkalle 42, DK-2100 Copenhagen^17^. All unique *in-silico* extracted 14 amino acid overlapping 16-mer peptides were randomly split into 6 evenly sized groups and synthesized in 6 equally sized sectors on the array. Each peptide was synthesized in a 20×20 µm^2^ field, separated from other fields by 10 µm on all sides. As a visual positive control, the amphipathic helix Hecate (peptide sequence: LKKALKKLKKALKKAL) was synthesized in the 17 peptide fields encompassing each of the corners of the 6 sectors on the array. Hecate and other controls, including 510 blank fields, were also spread randomly across the array. All peptides were synthesized onto the array anchored at the C-terminal with a KSGG linker. The array was blocked using 5% BSA in PBS with 2 mM DTT for 1h on a shaker at 120 rpm and then washed 3 times in PBS for 20 min. The array was incubated with DiD-liposomes diluted 1:100 in PBS (pH 7.4) for 1h at RT. Subsequently, the array was washed twice in PBS for 10 min and an additional time for 20 min. Following 3×30 second washes with 0.1% Tween-20 in PBS (pH 7.4) the array was dried by centrifugation in a table centrifuge (30 seconds) and scanned at 1-µm resolution with an InnoScan 900 laser-scanner (INNOPSYS) with an excitation wavelength of 635 nm and gain setting 6 with the high-power setting.

#### Array mapping, correction and normalization

A grid of size 177 × 210 was manually fit to each of the six sectors of the array image to identify individual fields (using in-house software by Schafer-N Aps). Fields containing clear signal artifacts were labeled and given a signal intensity of NA. The average signal intensity, sector identifier and location of each field within the sector was extracted and mapped back to the synthesis file to identify the corresponding peptide sequence. We corrected for signal gradients on the array by multiplying the signals with a mask created by kernel smoothing using the Nadaraya/Watson normalization of the kernels to handle missing data (Figure S3C). The correction made the data indistinguishable from gradients of arrays with evenly distributed signals as produced by bootstrap sampling (Figure S3D-E). Corrected signals were normalized to the average signal intensity of blank fields and the average signal intensity of control fields containing Hecate.

#### Protein reconstruction and statistical testing

To create binding profiles for each terminal, normalized intensities for the corresponding peptide fields were plotted in sequence order (Figure S4A). Missing values were imputed from the distribution of mean normalized intensities from every terminal (Figure S5A). To correct for peptide position we determined the fluorescence intensity contribution of each amino acid pair in the sequence by calculating the average fluorescence intensity of peptide fields containing the corresponding amino acids (Figure S4B-C). To remove noise we used the fast Fourier transformation to remove 80% of all frequencies (Figure S4D-F). Local minima and maxima were determined for all terminals and peaks were separated if a local minima was at least 50% lower than the previous maxima (Figure S6A and D). To test the significance of the resulting peaks we assembled 10,000 mock terminals for each of the lengths of all true TMP terminals by bootstrap sampling random fluorescence intensities from peptide fields on the array. By calculating the maximum binding scores for peaks in the mock terminals of every length, we created background distributions of the maximum scores expected by random for terminals and peaks of all lengths (Figure S5B-E). We fit power-normal probability curves to all distributions and calculated the probability of observing the peaks from true reconstructed termini (Figure S5). A significance criterion of 0.05 was used to classify peaks into binding and a significance criterion of 0.95 for non-binding, and a score corresponding to the area under the curve was assigned to each peak (AUC score).

#### Calculation of substitution scores

The pre-computed phylogenetic tree for 60 vertebrate genomes available in the Ensembl genome browser was converted into a rooted and noded tree. Orthologs for each protein were retrieved from rest.ensembl.org (2022-03-10) and a single ortholog for each species with the highest sequence coverage with the target sequence was selected. Sequence coverage for each species ortholog was calculated after sequence alignment using the ClustalOmega algorithm with a BLOSUM80 substitution matrix (https://ftp.ncbi.nih.gov/blast/matrices/) implemented in the msa package in R^78^. Subsequently a multiple sequence alignment was run with all selected sequences as described above. Sequences with more than 50% mismatches were discarded. For each position in the multiple alignment, the amino acid for each species was used to estimate the ancestral character states of the fixed phylogenetic tree. The amino acid probability for each node was calculated as a maximum likelihood estimation with an equal-rates model^79^. An amino acid state was determined for each node by considering the product of the amino acid probability and the substitution cost. The substitution cost was derived from a modified BLOSUM80 substitution matrix with values ranging from 0-1, 1 representing self-substitution and 0 as the theoretical maximum cost as done by Chang et al.^50^. Next, a weighted substitution score was calculated as the sum of the substitution cost at each node of the tree. A relative weighted substitution score was calculated by dividing the weighted substitution score with the mean weighted substitution score of each amino acid in the multiple alignment. Calculations and data retrieval were performed using the programming language R (v4.0.2).

#### Encoding and clustering of Amphipathic helices

For each binding site classified as an amphipathic helix, the 11 cassette with the maximum <µH> was determined, as done by Gautier et al.^75^. The position of each amino acid in the helical wheel projection with respect to the <µH>-vector was extracted and the resulting positions were binned into 18 bins. Each amino acid was assigned a hydrophobicity score according to its octanol/water partition^80^. Amino acid hydrophobicities in the same bin were summed. The resulting vectors of length 18 were joined into a matrix with 18 columns and principal components were calculated. The four principal components explaining most of the variance were used for subsequent K-means clustering and UMAP projection.

#### Molecular biology

Coding sequences for amphipathic helices were synthesized and cloned in frame into a previously described pcDNA3 Flag-Tac construct^81^ using HindIII and XbaI sites, ligated using T4 DNA ligase, and transformed into XL2-Blue. The amino acid sequence of Tac is largely equivalent to the Interleukin-2 receptor subunit ɑ, with a Flag-tag (DYKDDDDK) sequence inserted downstream of the signal peptide sequence and a trailing QASS sequence at the C-terminus. The C-terminal serine was replaced with amphipathic helices using primers listed in the key resources table.

#### Transient transfection for internalization assay

A day prior to transfection, cells were seeded on poly-L-ornithine treated coverslips in TPP polystyrene 12-well plates at a density of 2.5×10^5^ cells/well. The following day cells were transfected using Lipofectamine 2000 and Opti-MEM. The manufacturer’s protocol was followed with a 3:1 Lipofectamine to DNA ratio. A total of 1 µg/well was used and cells were incubated with transfection mix in Opti-MEM for 5 hours. Opti-MEM was replaced with growth media, and cells were used for experiments 48 hours after transfection.

#### Internalization assay

On the day of the experiment, cells were incubated with primary anti-FLAG M1 antibody (1 µg/mL in serum-free media, for 1 hour at 4^°^C). Cells were then exposed to pre-warmed (37^°^C) growth media containing 25 µM Monensin and kept at 37^°^C for 30 minutes. Cells were fixed in a 4% PFA solution in PBS for 10 minutes and subsequently washed with PBS. Next, cells were passivated using a blocking buffer (5% GS in PBS) and incubated for 20 minutes at 25^°^C. Membrane-bound M1 antibody was labeled using Alexa Fluor^®^ 488 conjugated goat anti-mouse secondary antibody (1:200 in 5% GS, 1 hour, 25^°^C). Cells were fixed again in 4% PFA in PBS for 10 minutes at 4^°^C, washed in PBS and permeabilized with 0.2% saponin and 5% GS in PBS (30 minutes at 25^°^C). Next, cells were incubated with Alexa Fluor^®^ 647 conjugated secondary goat anti-mouse antibody (1:500, 0.2% saponin and 5% GS, 30 minutes 25^°^C). Cells were washed twice in PBS and incubated with Hoechst stain (1:2000) for 2 minutes. Following three washes with PBS coverslips were mounted on glass slides with ProLong Gold Antifade Mountant and left to dry in the dark.

#### Confocal imaging

Confocal z-stack images were acquired using an inverted Zeiss Axio Observer Z1 microscope coupled to a Hamamatsu Orca Fusion Yokogawa spinning disk and a Plan-Apochromat oil immersion objective with a 63x magnification and NA of 1.4 (DIC III) (Carl Zeiss, Oberkochen, Germany). Samples were imaged in 1024×1024 resolution with a bit-depth of 16 bit. Hoechst was excited with a 405nm Diode and emitted fluorescence was filtered using a 450/50 band pass filter. Alexa Fluor^®^ 488 was excited with a 488-nm Argon laser and emitted fluorescence was filtered using a 525/50 band pass filter. Alexa Fluor^®^ 568 was excited with a 561-nm HeNe laser and emitted fluorescence was filtered using a 629/62 bandpass filter. Alexa Fluor^®^ 647 was excited with a 635-nm HeNe laser and emitted fluorescence was filtered using a 690/50 bandpass filter. Internalization experiments were imaged using Hoechst, Alexa Fluor^®^ 488 and Alexa Fluor^®^ 647, while colocalization experiments were done using Hoechst, Alexa Fluor^®^ 488 and Alexa Fluor^®^ 568. Channels were imaged separately.

#### Z-stack image processing

Image processing was performed using FIJI ImageJ. Regions of interest were selected to accommodate entire cells throughout the stack, as inspected through the green channel only. All regions of interest have been saved for later inspection. For the internalization assay, an in-house processing pipeline was developed using an ImageJ macro and a post-processing script written in R. In short, thresholds were set to exclude background noise, kept constant throughout independent experiments, and all images were masked. The average intensity for each region of interest was quantified independently for each stack in each channel, summed, and saved as comma-separated files. The post-processing script was used to calculate overall intensity ratios for each cell. All set thresholds have been saved for later inspection.

#### Colocalization analysis

Confocal images were obtained on the inverted Zeiss Axio Observer Z1 spinning disk as described. Subsequent image processing was performed using FIJI ImageJ. Regions of interest were selected as described in Z-stack image processing and saved for later inspection. The images were background subtracted and analyzed using the ImageJ BIOP JACoP plugin with manual thresholds to acquire Pearson’s correlation coefficient and Manders’ colocalization coefficients M1 and M2. Analysis is kept as an ImageJ script for later inspection. Data was plotted and statistical tests were performed in GraphPad prism.

### QUANTIFICATION AND STATISTICAL ANALYSIS

If not otherwise specified, statistical analysis was performed in R v4.3.1 (IDE: RStudio v2023.06.1, posit.co) using the **stats** (v4.3.1) and **gamlss** (v5.4-20) packages. Heatmaps were produced using the **ComplexHeatmap** (v2.16.0) package^82^. Protein structures were produced using **r3dmol** (v0.2.0). Sequence alignments were performed using the **msa** package (v1.32.0). Sequence similarity matrices were calculated using *ParSeqSim* in the **protr** package (v1.7.0). Kernel density estimates were performed using the **MASS** package (v7.3.60). PANTHER overrepresentation analysis was done using pantherdb.org via the **rbioapi** package (v0.8.2). Graphs were produced using **ggplot2** (v3.5.0) and **ggprism** (v1.0.5). Functional analysis was performed in Cytoscape (v3.9.1) with ClueGO (v2.5.10)^83^. Kruskal-Wallis tests with Dunn’s correction for multiple comparisons in Figure 4 and Figure 7 were performed in Prism v.10.1.1 and v.10.2.1, respectively. For sequence motif discovery, we used the STREME algorithm^84^ to identify statistically significant motifs in non-amphipathic helix binding sites (p<0.05, <µH> < mean of non-binding, AlphaFold plDDT<50) against the non-binding background (p>0.95). Logoplot for Figure 7F was created with STREME.

### ADDITIONAL RESOURCES

For non-programmatic access, the dataset has been made available as a shiny app: MemRIDRdb

